# Designing a robust whole-cell biosensor for detection of toxic metals using intein splicing inhibition of *Mycobacterium tuberculosis* SufB protein

**DOI:** 10.1101/2025.09.27.678972

**Authors:** Ashwaria Mehra, Ananya Nanda, Sasmita Nayak

## Abstract

Disruption of the natural geochemical cycle by human activities has led to bioaccumulation of metals, posing a global health threat. Hence, there is a pressing need for simple, sensitive, yet eco-friendly biosensor setups to monitor metal contamination in the environment. Existing biosensors are limited by poor efficiency, stability issues, and complex instrumentation requiring skilled operators. To address these caveats, current study explores how intein-mediated protein splicing, a spontaneous post-translational process, can be adapted for metal-biosensing by coupling metal-dependent splicing inhibition to viability loss of native microbial cells. Toxic metal ions like Cd^2₊^ and Hg^2₊^ attenuated the splicing activity of *Mtb* SufB precursor protein over a concentration range of 25 µM to 2 mM, while Pb^2₊^ and Cr^3₊^ failed to do so. An innovative biosensor platform was designed for colorimetric detection of metal ions via simple Alamar Blue assay, where attenuated *Mtb* strain (H37Ra) served as the indicator cells. Metal-induced SufB splicing inhibition led to loss of viability of H37Ra cells, while addition of metal-specific chelators reversed the effect. Multiplexing ability was evaluated by including known splicing inhibitors like Cu^2₊^, Zn^2₊^, and Pt^4₊^ over various concentration range alongside Cd^2₊^ and Hg^2₊^. The simple 96-well plate format enables multiplexed qualitative metal detection, while colorimetric absorbance measurement ensures metal quantification. The designed biosensor offers low-cost, user-friendly, and sensitive assay for high-throughput metal detection, utilizing whole-cell native organisms carrying metal-sensing precursor protein. Thus, this approach can be implemented in standard biological laboratories for robust metal screening process in environmental and industrial effluents.

## Introduction

Heavy metal contamination has emerged as one of the most persistent environmental and public health challenges of the modern era. Over 35 metals have been identified in their ore forms within the Earth’s crust, many of which are essential nutrients. Among these, 23 metals are classified as heavy metals due to their high density (>5 g/cm^³^) and atomic weights (>40.0), indicating their potential for environmental toxicity (1–3). Human activities have significantly disrupted natural geochemical cycles, causing bioaccumulation and biomagnification through contaminated food, water, or direct exposure, thereby posing serious health risks (3–7).

Current analytical methods, such as electrothermal atomic absorption spectrometry (ETAAS), inductively coupled plasma-optical emission spectrometry (ICP-OES), high-pressure liquid chromatography combined with ICP-mass spectrometry (HPLC-ICP-MS), hydride generation atomic absorption spectrometry (HGAAS), and X-ray fluorescence spectrometry (XRF), offer excellent sensitivity for detecting metals. Nonetheless, these methods involve complex instrumentation, require skilled operators, generate hazardous waste, contributing further to environmental pollution (8–11). To overcome these limitations, biosensors have emerged as a promising alternative offering affordability, and long-term operational stability (8). According to the International Union of Pure and Applied Chemistry (IUPAC), biosensors integrate biological recognition elements directly with transducers to provide precise quantitative data (12, 13). Although enzyme-based electrochemical biosensors have been widely explored, their broader implementation is limited due to poor electron transfer efficiency, and instability, limiting their commercial reusability (14–20). In comparison, whole-cell biosensors present a more practicable approach by using live cell sensing units, with advantages such as stable signal generation, and scalability. For instance, Chouteau et al. synthesized BSA encapsulated microalgae as bioreceptors to detect metals (Cd^2₊^, Zn^2₊^, and Pb^2₊^) (16). However, the sensitivity of detection was affected due to variability in algal loading and distribution within BSA membrane. In another work, transformed protozoan cells were used as whole-cell biosensor, linking the metallothionein promoter-based metal detection to luciferase reporter system (21). Despite the innovation, signal variability compromised the specificity and sensitivity of heavy metal detection. In another study, Kim and colleagues engineered *E. coli* cells carrying plasmids with metal sensing promoters to detect metals such as Cu^2₊^, Zn^2₊^, and Cd^2₊^ (22). However, the Cu^2₊^ and Cd^2₊^ sensing strains were not selective and could also detect other metals. Further, detection of Cu^2₊^ was hindered by the active transport of Cu^2₊^ ions out of the cytoplasm by the endogenous CopA protein, suppressing reporter gene expression. Other challenges with whole-cell biosensors include declining cell viability causing delayed response time, leaky gene expression inducing false positive signals, and ecological threats of using genetically modified organisms (23–29). Addressing these shortcomings, intein-mediated protein engineering techniques open up new avenues for creating innovative cell-based sensors (30–33).

Inteins are protein splicing elements within precursor proteins that can self-excise via multi-step nucleophilic displacement reactions, to generate active proteins (Supplemental Fig. S1) (34, 35). This is a post-translational process that does not require energy sources or cofactors, hence inteins can be seamlessly integrated into biosensor units minimizing interference with sensor cell functionality (30). Furthermore, intein splicing in precursor proteins can be regulated via conditional protein splicing (CPS), in presence of small molecules such as metal ions and metal-based nanoparticles (Table 1) (36–42). For instance, a recent study from our group has examined the splicing and N-terminal cleavage activity of *Mycobacterium tuberculosis* (*Mtb*) SufB precursor protein in the presence of metal ions such as Cu^2₊^, Zn^2₊^, Pt^4₊^, and Fe^3₊^/Fe^2₊^ (43). Prior studies have also reported Zn^2₊^ mediated splicing regulation of PI-*Sce* VMA intein, *Mtb* RecA intein, and *Ssp* PCC6803 DnaE split intein (44, 45). Effects of metal ions, such as Cu^₊/2₊^ and CisPt, have been evaluated on *Mtb* RecA, *Cryptococcus neoformans* Prp8, and *Mycobacterium smegmatis* DnaBi1 intein splicing (38, 39, 44, 46–51).

**Table 1:**
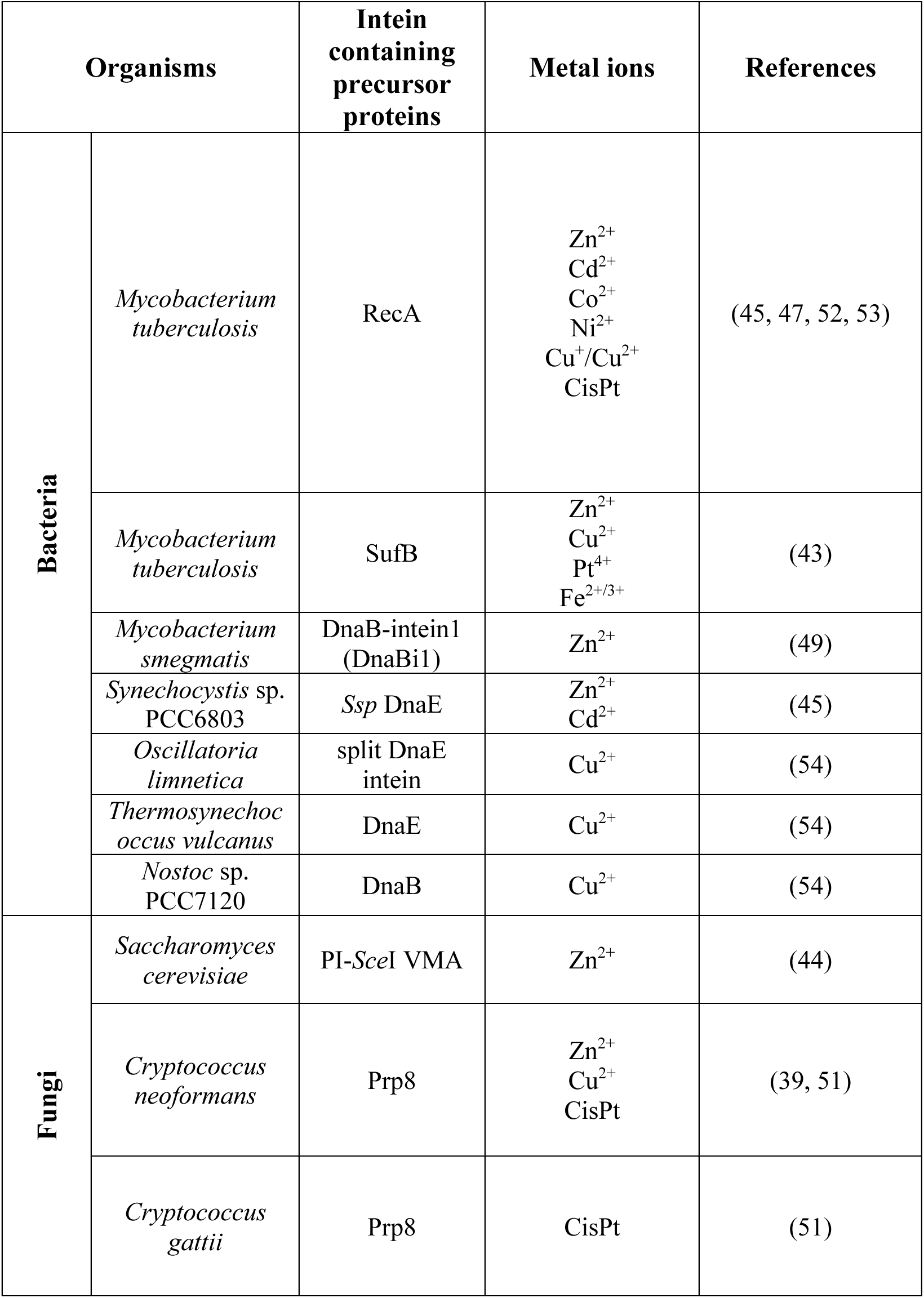
Overview of intein-harboring organisms and reported modulation of splicing activity by metal ions.

Conditional splicing activity of intein-containing proteins has been used to (30, 55–58) design biosensors to detect protein∼protein interaction in living cells via split GFP (52) or Luciferase reporter system (53–55). Although several fundamental studies have reported splicing regulation of precursor protein in presence of oxidative/nitrosative stresses (56), metal ions (43), nanoparticles (40) correlating to pathogen survival, a versatile and native whole cell intein-based biosensor, especially for toxic metal ion detection, is yet to be elucidated.

However, concerns pertaining to the functionality and usability of the intein-based metal-sensing biosensor remains, such as: 1) If an eco-friendly whole cell biosensor system can be designed using native cells without any genetic modification; 2) Whether the designed biosensor can offer an user-friendly and low-cost approach?; 3) If the working protocol can be simple, yet effective for a specific, and sensitive metal detection?; And finally, 4) Whether this approach can be applied for a multiplexed metal screening process, for qualitative and quantitative detection of different metal ions? Current research explores how toxic metal ions, like cadmium (Cd^2₊^), mercury (Hg^2₊^), chromium (Cr^3₊^), and lead (Pb^2₊^), affect the splicing and N-terminal cleavage reactions of *Mtb* SufB precursor protein. The fundamental knowledge gained from these results was applied to develop a biosensor platform to detect toxic metals, using native mycobacterial cells in a 96-well plate format. Splicing regulation in metal-sensitive precursor protein *Mtb* SufB induced alteration in mycobacterial cell viability, that indicated the presence of metal ions.

*Mtb* SUF protein complex constitutes an unique pathway for Fe-S cluster assembly in mycobacteria, and its activity is dependent on the splicing of SufB precursor, producing functional SufB protein (60). Thus, splicing inhibition which blocks the generation of active SufB, can retard mycobacterial growth and viability by impairing Fe-S cluster biogenesis (61). In the current work, *in vitro* studies demonstrated significant inhibition of *Mtb* SufB precursor protein splicing in presence of toxic metal ions such as Cd^2₊^, and Hg^2₊^ over a concentration range of 25 µM to 2 mM. Further, time-dependent assay revealed that Cd^2₊^ and Hg^2₊^ inhibit *Mtb* SufB intein splicing as early as 10 minutes following exposure to the metal ions. Building on this observation, a biosensor platform was designed using native H37Ra (an attenuated *Mtb* strain) as the metal-indicator cells. Alamar Blue assay performed in a 96-well microtiter plate, detected loss of cell viability in SufB-containing H37Ra cells in presence of toxic metal ions, Cd^2₊^ and Hg^2₊^. In a control study, *Mycobacterium smegmatis* (*M. sm*), which expresses an intein-less SufB protein, persisted as healthy, viable cells under similar experimental conditions. These results highlight the sensitivity of H37Ra-based biosensor platform to detect toxic metal ions over a concentration range of 25 µM to 2 mM. The specificity of the designed biosensor was assessed by adding respective metal chelators, which protected mycobacterial cell viability, by blocking the splicing-inhibitory activity of metal ions. Moreover, multiplexing and quantitative potential of the biosensor platform was validated by using various concentrations of known splicing inhibitors [Zn^2₊^, Cu^2₊^, and Pt^4₊^ (43)] alongside Cd^2₊^ and Hg^2₊^. The high throughput design enables simultaneous testing of many samples for a qualitative metal detection, while absorbance measurement from the colorimetric Alamar Blue Assay ensures metal quantification extrapolated to a standard plot. Thus, this work opens new avenues for developing a reliable and sensitive biosensor platform using whole-cell mycobacteria as biological indicator for toxic metal ions as well as trace metal elements (TME) in the environmental samples. The efficacy of this system lies in its user-friendly design, specificity, and sensitivity, enabling multiplexed metal detection by non-genetically modified whole cell microbes. Consequently, the working principle can be easily optimised and employed in standard biological laboratories equipped with organisms harboring other metal-sensitive intein-carrying precursor proteins.

## Experimental Procedure

### Experimental Design and Statistical Rationale

Current study was designed to evaluate intein splicing-based whole-cell mycobacterial biosensor, where splicing inhibition in the metal-sensing SufB precursor protein, reduced mycobacterial viability. During *in vitro* refolding assay, purified denatured *Mtb* SufB precursor and *Mtb* SufB splicing inactive (SI) double mutant (Cys1Ala/Asn359Ala) (splicing negative control) proteins were renatured in the absence and presence of CdCl₂, HgCl₂, CrCl₃, and PbCl₂ over the test concentration range (2.5 µM–2 mM). All the experiments were performed in triplicates (n=3) and the protein products from the Coomassie-stained SDS–PAGE gels were quantified by densitometric analysis using GelQuant.NET biochemical solutions software. The splicing and cleavage efficiencies were calculated and expressed as percentage values after baseline correction, using 0 hour splicing and cleavage value(s).

For evaluation of the metal biosensor, a simple Alamar Blue assay was conducted, where different mycobacterial cells [*Mtb* H37Ra cells and *Mycobacterium smegmatis* (SufB intein-less)] were incubated with varied concentrations of Cd^2₊^ (2.5 µM- 2 mM) and Hg^2₊^ (25 µM- 2 mM) in 96-well microtiter plate. In a parallel experimental setup, Cd^2₊^ and Hg^2₊^ specific metal chelators were added in a ratio of 1:1 (metal: chelator) to examine the specificity of the biosensor platform. For a quantitative evaluation, standard plots were generated by correlating the known concentrations of samples [Cd^2₊^ and Hg^2₊^] along the X-axis, to the respective absorbance values from the Alamar Blue assay on the Y-axis. Test metal concentrations in the unknown samples were interpolated from the standard curve to validate the biosensor efficacy. Multiplexing ability of the whole-cell biosensor was assessed using known splicing inhibitors ZnCl₂, CuCl₂, and PtCl₄, along with CdCl₂ and HgCl₂, under identical experimental conditions.

Statistical analysis for refolding assays was performed using one-way analysis of variance (ANOVA) to compare splicing and N-terminal cleavage efficiencies of metal treated samples against the untreated protein control, after baseline correction using 0 h values, and results were plotted using GraphPad Prism version 5.01 for Windows, GraphPad Software, SanDiego, CA, U.S.A. (www.graphpad.com). The data are presented as mean ± SEM. The statistical significance (p-value) between the control and experimental groups is denoted by * (p < 0.05), or ** (p < 0.01), or *** (p < 0.001), or ****(p<0.0001).

### Protein overexpression and purification (62–64)

Plasmids containing the genes for *Mtb* SufB precursor and *Mtb* SufB splicing inactive (SI) double mutant (Cys1Ala/Asn359Ala) (splicing negative control), both carrying an N-terminal 6X (His) tag, were borrowed from Belfort Lab, SUNY, Albany, USA)(62). The test plasmids were transformed into BL21(DE3) *E. coli* cells (NEB, C2527I), followed by IPTG (500 μM, sigma 367-93-1) induced protein over-expression. Thereafter, lysis buffer (20 mM sodium phosphate, 0.5 M NaCl, pH 7.4) was used to resuspend the bacterial cell pellet and cells were lysed via tip sonicator (Sonics vibra cell VCX-130). The cell lysate was centrifuged at 16500×g for 20 minutes at 4 °C, and the supernatant was discarded. The collected inclusion body (IB) material was then washed three times in lysis buffer via centrifugation (16500×g for 20 minutes at 4 °C). Finally, the IB material was solubilized for protein extraction using 8 M urea (Merck, 1084870500) buffer (20 mM sodium phosphate, 0.5 M NaCl, 8M urea, 10 mM of imidazole, pH 7.4 (MP–biochemicals-288-32-4) and subsequently centrifuged at 16,500×g for 20 minutes at 4 °C to collect the supernatant. Then, solubilized proteins were purified by Ni-NTA affinity column (Ni-NTA His trap, HP GE healthcare life sciences, 17524802) (38, 45, 58, 65, 66). Before applying the samples, the columns were equilibrated with binding buffer (20 mM sodium phosphate, 0.5 M NaCl, 8 M urea, 10 mM imidazole, pH 7.4). Following sample loading, the columns were washed multiple times (5 CV) using wash buffer (20 mM sodium phosphate, 0.5 M NaCl, 8 M urea, 40 mM imidazole, pH 7.4). Subsequently, the test proteins were eluted as purified fractions using elution buffer (20 mM sodium phosphate, 0.5 M NaCl, 8 M urea, 500 mM imidazole, pH 7.4). Protein quantification was performed using Bradford reagent.

### Refolding assay for Mtb SufB precursor protein

Denatured protein (2.5 µM) in elution buffer was renatured in refolding buffer (20 mM sodium phosphate, 0.5 M NaCl, 0.5 M L-Arginine) containing 2 mM TCEP-HCl (sigma-51805-45-9) for 4 hours at 25 °C in the absence and presence of toxic metal ions (45, 65, 66). Gradient assay for toxic metal ions [CdCl_2_ (SRL-99643), HgCl_2_ (SRL-25699), CrCl_3_ (CDH-025702), PbCl_2_ (SRL-69789)] was performed using varying concentrations of toxic metal ions (2.5 µM to 2 mM). 0h sample was collected before initiation of refolding, and the reaction was abruptly blocked by the addition of loading dye (0.1% bromophenol blue, 50% glycerol, 5% β-mercaptoethanol, 10% SDS, tris 6.8), followed by rapid freezing at −20 °C. After 4h of protein renaturation, the protein samples were resolved through 4∼10% gradient SDS-PAGE, followed by staining with Coomassie brilliant blue R-250 (93473, SRL), and densitometric analysis was performed using GelQuant.NET biochemical solutions software. The splicing and cleavage efficiencies were calculated using the following formulae:

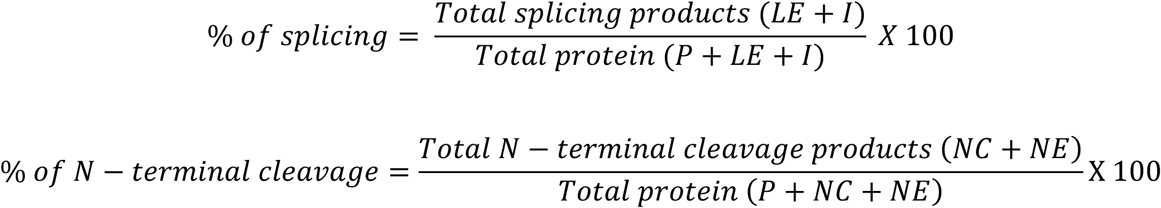

[P: precursor protein, LE: Ligated exteins, I: Intein, NC: N-terminal cleavage product, NE: N-extein]

Baseline correction was done using the 0 hours splicing and cleavage value(s). The results were analyzed using one-way ANOVA and plotted using GraphPad Prism version 8 for Windows, GraphPad Software, San Diego, California, USA, www.graphpad.com.

### Immunoblotting to identify splicing and N-terminal cleavage protein products(43)

Protein immunodetection was conducted to identify of splicing and N-terminal cleavage reaction products, using anti-6X(His) tag antibodies (Invitrogen, LOT 1902132). After resolution through 4∼10% gradient SDS-PAGE, the protein products were transferred onto a PVDF membrane (Millipore, IPVH 00010) at 50V for 2 hours at 4 °C. The blot was then blocked with 5% skim milk for 2 hours at room temperature, followed by two washes with 1X TBST. It was incubated with primary anti-6X(His) tag (abgenex-32-6116) at a dilution of 1:5000 at 4 °C for 16 hours. After washing with 1X TBST, the blot was treated with a secondary antibody (anti-mouse; abgenex 11-301) at a dilution of 1:8000 for 2 hours at 37 °C. The N-extein band was detected using a lower dilution of the primary antibody (1:2500) and the secondary antibody (1:6000). Finally, the blot was developed using Enhanced Chemiluminescence substrate (Abcam, ab65628) to visualize the protein bands, and images were captured using in-house ImageQuant^TM^ LAS 500 facility (GE Healthcare). Blots were also developed, using 3, 3’-diaminobenzidine (DAB) (SRL-94524) for visualization of the protein bands.

### Spectroscopic and Elemental Profiling of Protein-Metal Interactions UV-Visible spectroscopy (67–69)

Denatured proteins were renatured *in vitro* in the presence and absence of toxic metal ions at their respective concentrations for 4 hours, as specified earlier. Subsequently, the treated samples were subjected to UV-visible spectroscopy (Cary 100 UV-Vis Agilent Technology) scanning in the visible region spanning 200-800nm. To rule out the interference from buffer-metal interactions, the refolding buffer (20 mM sodium phosphate, 0.5 M NaCl, 0.5 M L-Arginine) was incubated with metal ions (Cd^2₊^, Hg^2₊^, Cr^3₊^, and Pb^2₊^) at their respective concentrations under similar experimental conditions. This was used as a blank for baseline correction and was scanned alongside each sample. Absorbance versus wavelength plots were generated and analyzed using Origin (Pro) 8.5 software (Version 8.5, OriginLab Corporation, Northampton, MA, USA) to interpret the results.

### Tryptophan Fluorescence Spectroscopy

The purified denatured protein (2.5 µM) was allowed to renature in the presence and absence of metals ions (25 μM CdCl_2_, 500 μM HgCl_2_, 2 mM CrCl_3_, and 2 mM PbCl_2_) under identical experimental conditions. Following 4 hours of refolding, the test samples were analyzed via Fluorescence spectrometry (Cary Eclipse, Agilent Technologies) over a wavelength range of 300-600 nm and at an excitation wavelength of 280 nm (70). To rule out the buffer-metal interactions, the refolding buffer (20 mM sodium phosphate, 0.5 M NaCl, 0.5 M Arginine) was incubated with metal ions at their respective concentrations at 25 °C for 4 hours and then scanned alongside each treated sample. This served as the blank for baseline correction. The tryptophan fluorescence spectra for the test samples were recorded and analyzed by plotting intensity vs. wavelength using Origin (Pro) 8.5, Version 8.5, from OriginLab Corporation, Northampton, MA, USA.

### Inductively coupled plasma-optical emission spectroscopy (ICP-OES) analysis

Following *in vitro* refolding of *Mtb* SufB precursor in the presence of test metal ions, the protein products were resolved through 4∼10% SDS-PAGE. The resulting protein bands were visualized with Coomassie brilliant blue R-250. Subsequently, the gel bands for unspliced precursor protein, ligated extein, and intein were excised from the gel for further analysis. Each gel band was then treated with concentrated H_2_SO_4_ and 30% H_2_O_2_. Complete dissolution of each sample was achieved by incubating the gel pieces in a 3:1 mixture of concentrated HCl and concentrated HNO_3_ at 80°C for 4 hours (71, 72). The final volume of the digested solution was adjusted to 50 ml with deionized water and then analyzed by ICP-OES (Perkin-Elner, optima 8300). Prior to analyzing the test samples, respective toxic metal standards were calibrated in the system, and the concentration of bound metal ions was quantified from the standard curve.

### H37Ra-based whole-cell toxic metal biosensor

Different mycobacterial cells [*Mycobacterium tuberculosis* H37Ra strain (ATCC 25177), *Mycobacterium smegmatis* (negative control for SufB intein-less micro-organism, ATCC 700084)] were cultured in 7H9 (Middlebrook 7H9 broth medium, M198) supplemented with OADC (Oleic acid, dextrose, and catalase, Himedia-FD018) and allowed to grow separately for 2-3 weeks at 37 ^0^C. CFU calculations for individual cultures were made by matching the turbidity of the inoculated mycobacterial culture with the McFarland standard (Himedia-R092) against a black background (73). Different bacterial cells were diluted in sterile saline to a concentration of 10^8^ CFU/ml. H37Ra cells were incubated with varied concentrations of Cd^2₊^ (2.5 µM- 2 mM) and Hg^2₊^ (25 µM- 2 mM) in 96-well microtiter plate. Since Cr^3₊^ and Pb^2₊^ over a test concentration range of 25 µM- 2 mM did not exert any effect on *Mtb* SufB splicing, these metal ions were not tested via the designed biosensor platform. The plates were sealed and incubated at 37 °C for 4 days. Then, 10% (v/v) solution of Alamar Blue reagent (SRL-42650) was added to each well, followed by incubation at 37 °C for 12 hrs.(74). Untreated mycobacterial cells grown in 7H9 (supplemented with OADC) media were used as a negative control, and Alamar blue reagent alone in 7H9 (supplemented with OADC) media served as a positive control. SufB intein-less organism, *Mycobacterium smegmatis,* was also evaluated under similar conditions and treated as an additional experimental control. After 12 hours of incubation, varied color changes were noted in different wells, where blue color indicated dead cells with no viable cell growth, and pink color correlated to metabolically active live cells. In a similar experimental setup, metal chelators specific for Cd^2₊^ (Edetate Calcium Disodium, ECD, Sigma-PHR1510) and Hg^2₊^ (meso-2,3-Dimercaptosuccinic acid, DMSA, Sigma-D7881) were added in a ratio of 1:1 (metal: chelator) to determine the specificity of the designed Biosensor platform.

### Quantitative evaluation of H37Ra-based whole cell toxic metal biosensor

Next, a standard plot was generated by measuring the absorbance of the respective samples at 570 nm, with 600 nm as a reference wavelength for quantitative analysis (74). A wide range of test metal samples (Cd^2₊^, Hg^2₊^) of known concentration were prepared and loaded onto the 96-well plate and Alamar Blue reagent was added under similar experimental conditions. The absorbance of test metal samples was calculated by subtracting the reference absorbance value at 600nm from the absorbance measured at 570nm [OD_test_ = OD_570_ – OD_600_]. Untreated mycobacterial cells grown in 7H9 (supplemented with OADC) media were used as a control for normalization. Using the normalized values, XY scattered plot was obtained where the X axis represented the known concentrations of the test metal ions and the Y axis denoted corresponding absorbance values. Next, to assess the applicability of the standard plot, four test samples (Unknown 1, Unknown 2, Unknown 3, and Unknown 4) containing varying concentrations of CdCl₂ and HgCl_₂_ were analyzed under identical experimental conditions. Absorbance values were extrapolated onto the standard curve to calculate the unknown metal ion concentrations, thereby validating the sensitivity and accuracy of the biosensor platform.

### Multiplexing evaluation of H37Ra-based whole cell metal biosensor

Bacterial cell suspension containing 10⁸ CFU/ml of *Mycobacterium tuberculosis* H37Ra were prepared in sterile saline. Cells were incubated with a varied concentration range of ZnCl_2_ (100 µM- 2 mM), CuCl_2_ (100 µM- 2 mM), PtCl_4_ (20 µM- 1 mM), CdCl_2_ (10 µM- 2 mM), and HgCl_2_ (25 µM- 2 mM), in 96-well microtiter plates. The plates were sealed and incubated at 37 °C for 4 days under static conditions. Then, 10% (v/v) solution of Alamar Blue reagent (SRL-42650) was added to each well, followed by incubation at 37 °C for 12 hrs. Untreated H37Ra cells grown in Middlebrook 7H9 supplemented with OADC media were used as a negative control, while Alamar blue reagent incubated in media alone served as a positive control. SufB intein-less organism, *Mycobacterium smegmatis,* was treated with test metal ions under similar conditions and used as negative experimental control to assess SufB intein-specific response. Post 12 hours of incubation, color change was noted in the wells, where blue color indicated dead cells with no viability, and pink color indicated healthy viable cells.

To further ascertain the specificity of the biosensor platform, respective metal-chelating agents were added in separate experimental setup, using a 1:1 molar ratio of metal ions to chelator. Cadmium-specific chelator Edetate Calcium Disodium (ECD; Sigma-PHR1510), mercury-specific chelator meso-2,3-Dimercaptosuccinic acid (DMSA; Sigma-D7881), and a broad-spectrum chelator Ethylenediaminetetraacetic acid (EDTA; HiMedia-MB011), were co-incubated with the respective metal ions under identical experimental conditions. Cell viability was evaluated using Alamar Blue reagent as described earlier.

### Statistical Analysis

Statistical analysis for refolding assays was performed using one-way analysis of variance (ANOVA) to compare splicing and N-terminal cleavage efficiencies of metal treated samples against the untreated protein control, after baseline correction using 0 h values, and results were plotted using GraphPad Prism version 5.01 for Windows, GraphPad Software, SanDiego, CA, U.S.A. (www.graphpad.com). The data are presented as mean ± SEM. The statistical significance (p-value) between the control and experimental groups is denoted by * (p < 0.05), or ** (p < 0.01), or *** (p < 0.001), or ****(p<0.0001).

## Results and Discussion

### Toxic metal ions inhibited Mtb SufB precursor protein splicing

Environmental concentrations of toxic metal ions such as cadmium (Cd^2₊^), mercury (Hg^2₊^), chromium (Cr^3₊^) and lead (Pb^2₊^) have been reported to vary widely. Cd^2₊^ concentrations typically ranges from 1 to 10 ppm (approximately 5 µM to 55 µM), Hg^2₊^ from 10 to 700 ppm (approximately 4 µM to 3 mM), Cr^3₊^ from 5 ppm-140 ppm (approximately 31 µM – 885 µM) and Pb^2₊^ levels from 10 to 50 ppm (approximately 36 µM to 180 µM) (75–90). Earlier studies have shown that metal ion concentrations between 40 µM to 2 mM effectively inhibit intein splicing (43, 91, 92). Built upon these observation, *in vitro* gradient assay was performed to evaluate the sensitivity of *Mtb* SufB precursor protein to a selected concentration range of toxic metals ions (2.5 µM to 2 mM) (Fig. 1, and Supplemental Fig. S2-S4). *Mtb* SufB precursor protein (95.98 KDa) follows a canonical intein splicing pathway yielding 40.2 KDa SufB intein and 55.7 KDa ligated exteins (functional SufB protein) as illustrated in Supplemental Fig. S1(62). N-terminal cleavage products (65.26 KDa N-terminal cleavage product, 29.9 KDa N-extein) and C-terminal cleavage products (70.12 KDa C-terminal cleavage product, 25.76 KDa C-extein) are the byproducts of *Mtb* SufB splicing reaction (62).

**Fig. 1.**
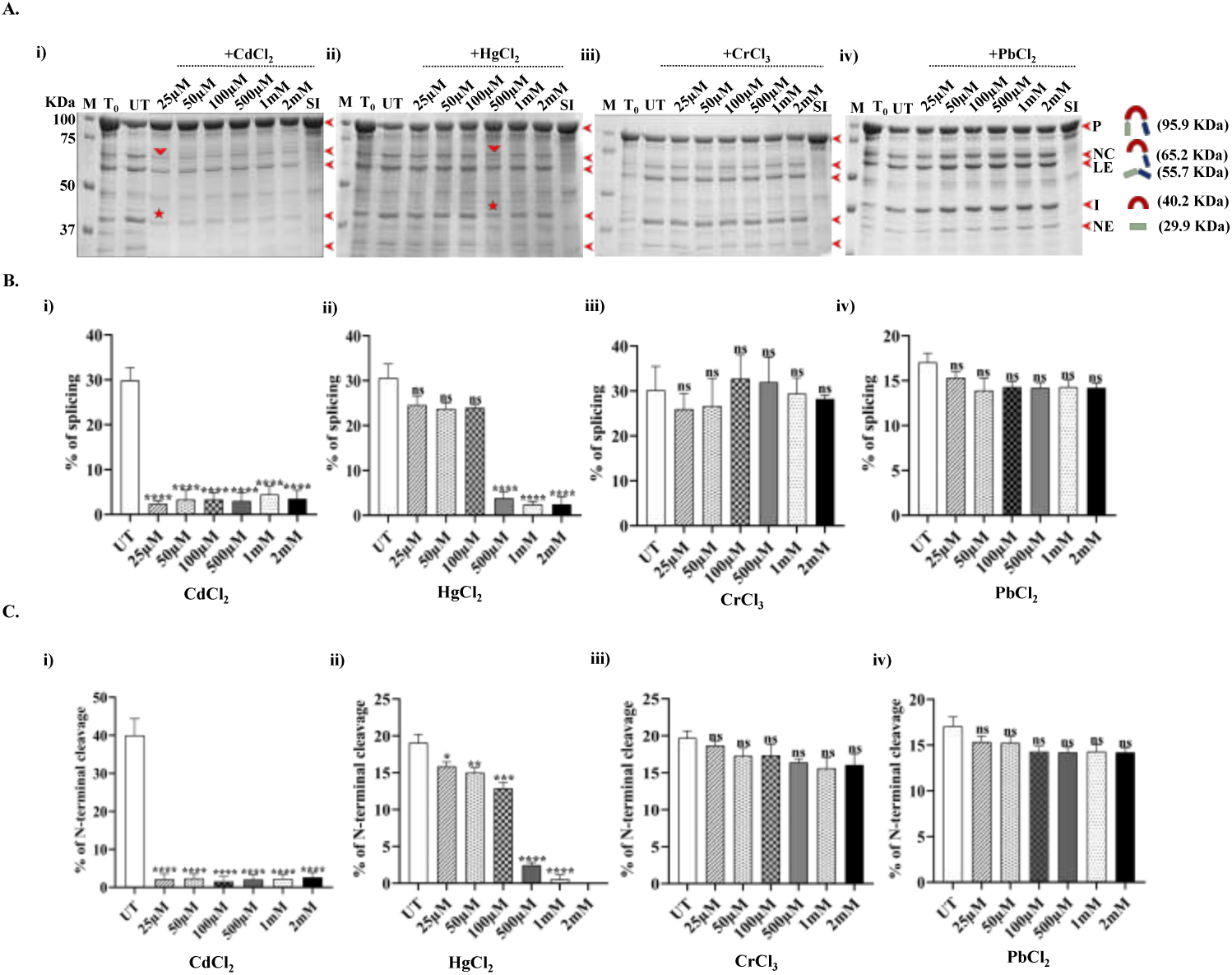
Gradient assay to determine the minimum effective concentration (MEC) of toxic metal ions on splicing and N-terminal cleavage reactions of *Mtb* SufB precursor protein. **(A)** Following *in vitro* refolding of *Mtb* SufB precursor protein in the presence of (i) CdCl_2,_ (ii) HgCl_2,_ (iii) CrCl_3,_ and (iv) PbCl_2_ at varied concentrations (25 µM-2 mM), resultant products were resolved through 4∼10% gradient SDS-PAGE. Lane 1: Protein ladder; Lane 2 (T_0_): splicing and N-terminal cleavage reactions at 0 h; Lane 3 (UT): splicing and N-terminal cleavage products of untreated protein sample; Lanes 4-9: splicing and N-terminal cleavage products resulting in the presence of different toxic metal ions at different concentrations (25 µM-2 mM). Lane 10 (SI): splicing inactive SufB double mutant (C1A/N359A) is used as a negative control for splicing. **(B) and (C)** Bar graphs demonstrating splicing and N-terminal cleavage efficiency of *Mtb* SufB precursor protein in the presence and absence of different toxic metal ions. A statistically significant difference in splicing and N-terminal cleavage reaction was observed for B (i) and C (i) CdCl_2,_ starting with 25 µM, and B (ii) and C (ii) HgCl_2_ at 500 µM concentrations, respectively. For CrCl_3_ [B (iii) and C (iii)], and PbCl_2_ [B (iv) and C (iv)], no significant difference in splicing and N-terminal cleavage reaction were observed between untreated and metal-treated samples. Graphs were plotted after densitometric analyses using GelQuant.Net biochemical solutions. All the experiments were performed in triplicate, and error bars represent (±1) SEM from 3 independent sets of experiments. The data shown in Fig. 1B, and 1C are extracted from Fig. 1A. M: Prestained protein molecular weight ladder, P: Precursor, NC: N-terminal cleavage product, LE: Ligated extein, I: intein, NE: N-extein.

Purified *Mtb* SufB precursor protein was refolded in presence of test metal ions and then resolved through 4∼10% gradient SDS PAGE (Fig. 1A). Cd^2₊^ at 25 µM, and Hg^2₊^ at 500 µM of minimum effective concentration (MEC) displayed significant attenuation of splicing and N-terminal cleavage reactions. There was a noticeable accumulation of SufB precursor protein, along with substantial reduction in the splicing products (ligated exteins and intein). Further, Cd^2₊^, and Hg^2₊^ at their respective MEC values and higher, induced a near-complete inhibition of N-terminal cleavage reaction and the products (N-terminal cleavage product and N-extein) (Fig. 1A). However, Cr^3₊^ and Pb^2₊^ failed to induce any effect on *Mtb* SufB splicing or cleavage reactions across the test concentration range (Fig. 1A). Resultant protein products were further confirmed via western blot using anti-6X(His) tag antibody (Supplemental Fig. S2). Densitometric analysis revealed a 12.52-fold (p<0.0001) reduction in splicing efficiency and an 18.7-fold (p<0.0001) reduction in N-terminal cleavage efficiency with 25 µM Cd^2₊^ [Fig. 1B (i) and C (i)]. Similarly, treatment with 500 µM Hg^2₊^ resulted in an 8-fold (p < 0.0001) reduction in splicing and a 7.8-fold (p < 0.0001) decrease in N-terminal cleavage efficiency [Fig. 1B (ii) and 1C (ii)]. No significant alterations in SufB splicing or N-terminal cleavage products were obtained in presence of Cr³₊ or Pb^2₊^ (Fig. 1B and 1C). All the test metal ions at a lower concentration range (2.5 µM- 10 µM) failed to exert any significant effect on the SufB splicing reactions (Supplemental Fig. S3 and S4). In order to rule out the possibility of precursor protein degradation by test metal ions, splicing inactive (SI) *Mtb* SufB double mutant (C1A/N359A) was refolded under similar experimental conditions (25 µM Cd^2₊^, 500 µM Hg^2₊^, 2 mM Cr³₊, and 2 mM Pb^2₊^). Since Cr³₊, and Pb^2₊^ showed no regulatory effects, highest test concentration range for both were included in the above study. SDS-PAGE and subsequent western blot analysis using anti-6X(His) tag antibodies confirmed the absence of protein degradation products (Supplemental Fig. S2B).

Further, a time-dependent splicing assay was performed to examine *Mtb* SufB splicing and cleavage activity over a period of 10 min- 4 hr., elucidating remarkable inhibitory roles of Cd^2₊^ (25 µM) and Hg^2₊^ (500 µM) as early as 10 minutes (Supplemental Fig. S5). Next, to evaluate the possible physical interaction between *Mtb* SufB protein and the metal ions, a series of experiments were conducted, as shown in the upcoming section. For the said interaction study, the MEC for Cd^2₊^, Hg^2₊^ (25 µM, 500 µM, respectively), which inhibited *Mtb* SufB splicing was selected. Since, Cr³₊, and Pb^2₊^ did not exhibit any such regulatory activity, highest value of the test concentration range (2 mM) was chosen to examine possible interaction with SufB protein.

## Protein∼ metal interaction study

### UV-Visible spectroscopic analysis

Biomolecules such as proteins exhibit a strong affinity for metal ions, which can critically influence their structure and function (93). Prior studies have demonstrated that toxic metal ions such as Cd^2₊^, Hg^2₊^, Cr³₊, and Pb^2₊^ can interact with calcium and zinc-binding sites in proteins, thereby influencing their biological activity (94, 95). UV-visible spectroscopic analysis was conducted to investigate the interaction between the *Mtb* SufB precursor protein and toxic metal ions under treated and untreated conditions. Absorbance spectra for *Mtb* SufB protein were recorded over 200-800 nm range at 25 °C (Supplemental Fig. S6). A characteristic absorbance peak near 280 nm was detected, primarily attributable to aromatic residues such as tryptophan and tyrosine (96, 97). Upon treatment with Cd^2₊^, an increase in absorbance intensity was noted, suggesting an alteration in conformation due to metal coordination [Supplemental Fig. S6 (i)]. Previous studies have mentioned that enhanced absorbance may result from the interaction between Cd^2₊^ and electron-donating groups such as amide, imidazole, and amine (68, 98, 99). Likewise, samples treated with Hg^2₊^ also exhibited an increase in absorbance intensity [Supplemental Fig. S6 (ii)]. This enhancement is likely attributed to the strong affinity of Hg^2₊^ for nucleophilic functional groups, particularly the sulfhydryl (–SH) groups of cysteine residues (100, 101). Although there were sharp increments in absorbance peak following interaction of Cr³₊and Pb^2₊^ with *Mtb* SufB, they failed to exert any effects on SufB splicing reactions as shown earlier (Fig. 1). Moreover, a strong buffer-Pb^2₊^ interaction was observed from the results, which may have weakened likely interaction with the protein. Although prior works have shown interaction of Cr³₊, and Pb^2₊^ with thiol, glutathione groups within protein structure including oxygen, nitrogen, and sulfur containing amino acids (7, 102, 103), current work could not detect any effects of these metal ions on *Mtb* SufB splicing activity [Supplemental Fig. S6 (iii and iv)]. To further substantiate such interactions, tryptophan fluorescence spectroscopy was performed, as explained in the forthcoming section.

### Tryptophan Fluorescence spectroscopy reveals metal-induced conformational changes in Mtb SufB

Tryptophan fluorescence spectroscopy is a widely utilized technique for monitoring alterations in protein conformation and assessing local structural dynamics upon interaction with ligands or macromolecules (104). Among the intrinsic fluorophores in proteins, aromatic amino acids—tryptophan (Trp), tyrosine (Tyr), and phenylalanine (Phe)—contribute to fluorescence, with tryptophan exhibiting the highest molar absorptivity (1614 cm⁻¹/M) and a characteristic emission peak in the range of 340–350 nm (105). The indole ring of Trp is primarily responsible for this emission and dominates the fluorescence profile over Tyr and Phe. Changes in the microenvironment of tryptophan residues, such as those caused by ligand binding, protein folding, or metal interaction, can lead to measurable shifts in fluorescence intensity and/or emission maxima (104, 106).

Altogether, 13 Trp residues are present, spanning the entire sequence of *Mtb* SufB precursor protein, with 5 Trp in N-extein, 5 Trp in Intein, and 3 Trp in the C-extein region (43). Upon treatment with Cd^2₊^ (25 µM), an increase in fluorescence intensity was observed [Supplemental Fig. S7 (i)], likely attributable to conformational rearrangements arising from interactions between Cd^2₊^ and functional groups such as amide, imidazole, and amine moieties within the protein matrix (68, 98, 99). These interactions may have led to reduced solvent exposure of Trp residues or stabilized a more rigid protein conformation, which enhanced fluorescence emission (68, 107). In contrast, Hg^2₊^ (500 µM) treated samples displayed a significant quenching of Trp fluorescence [Supplemental Fig. S7 (ii)]. This effect is consistent with previous studies indicating that mercury ions interact strongly with aromatic systems through cation–π interactions, particularly with the indole ring of tryptophan, thereby reducing fluorescence emission (108). Cr^3₊^ (2mM) exposure resulted in a slight increase in fluorescence intensity [Supplemental Fig. S7 (iii)], potentially due to the preferential binding of Cr³₊ to carboxyl groups, which may induce localized structural stabilization without extensive perturbation of the overall tertiary structure (109, 110). In case of Pb^2₊^ (2 mM), a decrease in fluorescence intensity was observed [Supplemental Fig. S7 (iv)]. This observation is supported by previous literature, where Pb^2₊^ interacts with hydroxy, amino, and sulfhydryl groups, leading to partial protein unfolding and increased solvent accessibility of Trp residues, thereby promoting fluorescence quenching (111, 112). Although Cr^3₊^, Pb^2₊^ have displayed possible alteration in protein conformation, they failed to regulate *Mtb* SufB splicing and cleavage reactions, likely due to lack of any effect on protein active site. Next, ICP-OES analysis was conducted to further ascertain potential interaction between test metal ions and protein backbone, as described in the following section.

### ICP-OES detects metal ions bound to SufB precursor and spliced out protein fractions

ICP-OES is a sensitive analytical technique capable of detecting trace metal concentrations by measuring the characteristic emission of photons from the excited atoms and ions as they return to their ground state (113). During *in vitro* protein refolding experiment, specific concentrations of Cd^2₊^ (25 µM = 4.583 µg/ml), Cd^2₊^ (500 µM = 91.66 µg/ml), Hg^₊2^ (500 µM = 135.76 µg/ml), Cr^₊3^ (2 mM = 361.72 µg/ml), and Pb^₊2^ (2 mM = 556.2 µg/ml) were added to the respective reaction mixture. ICP-OES analysis of the metal-bound protein fractions revealed detectable levels of Cd^2₊^ (2 ppb ∼ 0.002 µg/ml), Hg^2₊^ (5 ppb, ∼0.005 µg/ml), and Cr³₊ (80 ppb, ∼0.08 µg/ml) bound to the unspliced *Mtb* SufB precursor protein (Table 2). These findings indicate a direct and quantifiable physical interaction between the test metal ions and *Mtb* SufB precursor protein, which may underlie the observed inhibition of splicing and N-terminal cleavage reactions. Further, measurable interactions were observed between the test metal ions and various splicing products (ligated extein and intein) (Table 2). Cd^2₊^ addition to SufB protein sample at 25 µM concentration yielded negative results, while increasing the concentration to 500 µM led to positive data. At lower concentrations of Cd^2₊^ (25 µM), the metal content in the protein fraction was possibly below the instrument’s limit of detection (LOD) and limit of quantification (LOQ), resulting in negative readings. Although ICP-OES is capable of detecting metal concentrations at the ppb level, practical detection limits are influenced by matrix complexity and background interference, often resulting in quantification thresholds closer to the ppm range (114–116). Although 80 ppb (0.08 µg/ml) of Cr^3₊^ was bound to the unspliced precursor (Table 2), it failed to exert any visible effects on *Mtb* SufB splicing reactions (Fig. 1). No bound form of metal ions were detected in Pb^2₊^ treated samples, even at 2 mM concentration (Table 2), consistent with no visible effect on *Mtb* SufB splicing (Fig. 1).

**Table 2:** Quantitative elemental detection via ICP-OES analysis: Measurable concentration of metal ions (Cd^2₊^, Hg^2₊^, Cr^3₊,^ and Pb^2₊^) were identified bound to *Mtb* SufB precursor and splicing products. P: Precursor, LE: Ligated extein, I: Intein.

| Metal ions | Treated concentration | Concentration (µg/ml) | ICP-OES values (ppb) | Equivalent Concentration (µg/ml) |
| --- | --- | --- | --- | --- |
| Cadmium (Cd<sup>2+</sup>) | 25µM | 4.583 | P: -9<br>LE: -5<br>I: -8 | P: -0.009<br>LE: -0.005<br>I: -0.008 |
| Cadmium (Cd<sup>2+</sup>) | 500 µM | 91.66 | P: 2<br>LE: 18<br>I: 1 | P: 0.002<br>LE: 0.018<br>I: 0.001 |
| Mercury (Hg<sup>2+</sup>) | 500 µM | 135.76 | P: 5<br>LE: 3<br>I: 5 | P: 0.005<br>LE: 0.003<br>I: 0.005 |
| Chromium (Cr<sup>3+</sup>) | 2mM | 361.72 | P: 80<br>LE: 83<br>I: 90 | P: 0.08<br>LE: 0.083<br>I: 0.09 |
| Lead (Pb<sup>2+</sup>) | 2mM | 556.2 | P: -17<br>LE: -5<br>I: -73 | P: -0.017<br>LE: -0.005<br>I: -0.073 |

Collectively, the protein refolding assays, along with protein∼metal interaction studies, suggested possible interaction of Cd^2₊^ and Hg^2₊^ with *Mtb* SufB precursor protein, thereby attenuating the splicing and N-terminal cleavage efficacy. These mechanistic insights led to the development of a whole-cell biosensing platform utilizing H37Ra (*Mtb* attenuated strain) as the indicator cells to detect metal ions.

Fe-S clusters are essential cofactors for distinct metabolic pathways and cellular processes such as respiration, DNA replication and repair, and persistence under oxidative and nitrosative stress conditions (117–119). SUF complex offers the sole pathway for Fe-S cluster biogenesis in mycobacteria, and its activity relies on the generation of functional SufB protein via protein splicing (60, 61). Besides, there are earlier reports on loss of cellular viability as a result of SufB splicing inhibition within mycobacteria (40, 43, 56). Since, toxic metal ions like of Cd^2₊^ and Hg^2₊^ were shown to inhibit *Mtb* SufB splicing and N-terminal cleavage activity (Fig. 1), an attenuated strain of *Mtb* (H37Ra) was used as the whole-cell viability indicator for these metal ions.

### H37Ra-based whole-cell biosensor detects toxic metal ions

Cell-based biosensors are gaining interest as a screening platform for biologically active molecules due to their ability to monitor the cellular functions and interactions with targets in their natural environment (120, 121). Often, they are fabricated as genetically modified biosensors using native receptors or enzymes as molecular recognition elements in living cells (28). Current work explores the use of whole-cell mycobacteria (H37Ra)-based biosensor to detect toxic metal ions like Cd^2₊^ and Hg^2₊^. A 96-well plate assay was developed using Alamar Blue reagent, employing *Mtb* (H37Ra) cells as the sensing component, and *M. sm* cells as the SufB intein-less cellular control (56, 62, 122). *M. sm* carries three intein-bearing proteins: DnaB, which contains two inteins (DnaBi1 and DnaBi2), as well as GyrA and PhoH (49, 64). Although splicing regulation has been shown for DnaBi1 helicase, studies on the DnaBi2, PhoH, GyrA inteins splicing have not been reported yet. DnaBi1 splicing is regulated by zinc and oxidative stressors (49, 64); however, its helicase function can be compensated by other cellular helicases (123–125).

Alamar Blue reagent is a non-toxic, resazurin-based solution that functions as an indicator for cell viability (74, 126). The blue-colored non-fluorescent resazurin (7-hydroxy-3H-phenoxazin-3-one 10-oxide) gets reduced to resorufin (7-hydroxy-3H-phenoxazin-3-one, pink colored compound) by mitochondrial reductase in metabolically active cells (127). In a comparative study, non-viable H37Ra were visualized in presence of Cd^2₊^ and Hg^2₊^, whereas persistence of viable *M. sm* cells during the metal-interaction process highlighted the absence of metal sensing SufB precursor protein. Untreated H37Ra cells grown in 7H9 (supplemented with OADC) media, and Alamar Blue reagent alone in 7H9 (supplemented with OADC) media, served as the negative and positive controls for the study respectively.

Fig. 2 (i) is a schematic representation of 96-well plate-based biosensor platform containing *Mtb* H37Ra cells (indicator organisms) incubated with varied concentrations of Cd^2₊^ (2.5 µM -2 mM) and Hg^2₊^ (25 µM -2 mM). The test concentration range for the metal ions were chosen based on the positive results obtained during *in vitro* splicing analyses (Fig. 1, Supplemental Fig. S2-S4). For a baseline correction, H37Ra cells were grown in the absence of toxic metal ions (negative control). SufB intein-less, *M. sm* cells were used as additional experimental control. Unlike *Mtb* H37Ra cells, the growth and viability of the *M. sm* cells are independent of SufB intein splicing, and any loss of cell viability indicates metal toxicity. Fig. 2 (ii) shows that the indicator cells (H37Ra) incubated in the presence of lower concentration of Cd^2₊^ (2.5-10 µM) did not lose cell viability. All the cells were metabolically active and converted resazurin (blue) to resorufin (pink), since Cd^2₊^ at these concentration (2.5-10 µM) didn’t affect SufB splicing activity, generating active SufB protein [Supplemental Fig. S3 (i)]. H37Ra cells incubated in presence of 25 and 50 µM of Cd^2₊^, displayed purple color indicating a partial loss of cell viability, underscoring 25 µM as the minimum inhibitory concentration (MIC). Cells incubated with Cd^2₊^ at a concentration range of 100 µM- 2 mM, retained blue color, indicating complete loss of cell viability. These results correlated to inhibition of splicing and N-terminal activity of *Mtb* SufB in presence of Cd^2₊^ over a concentration of 25 µM- 2 mM (Fig. 1).

**Fig. 2.**
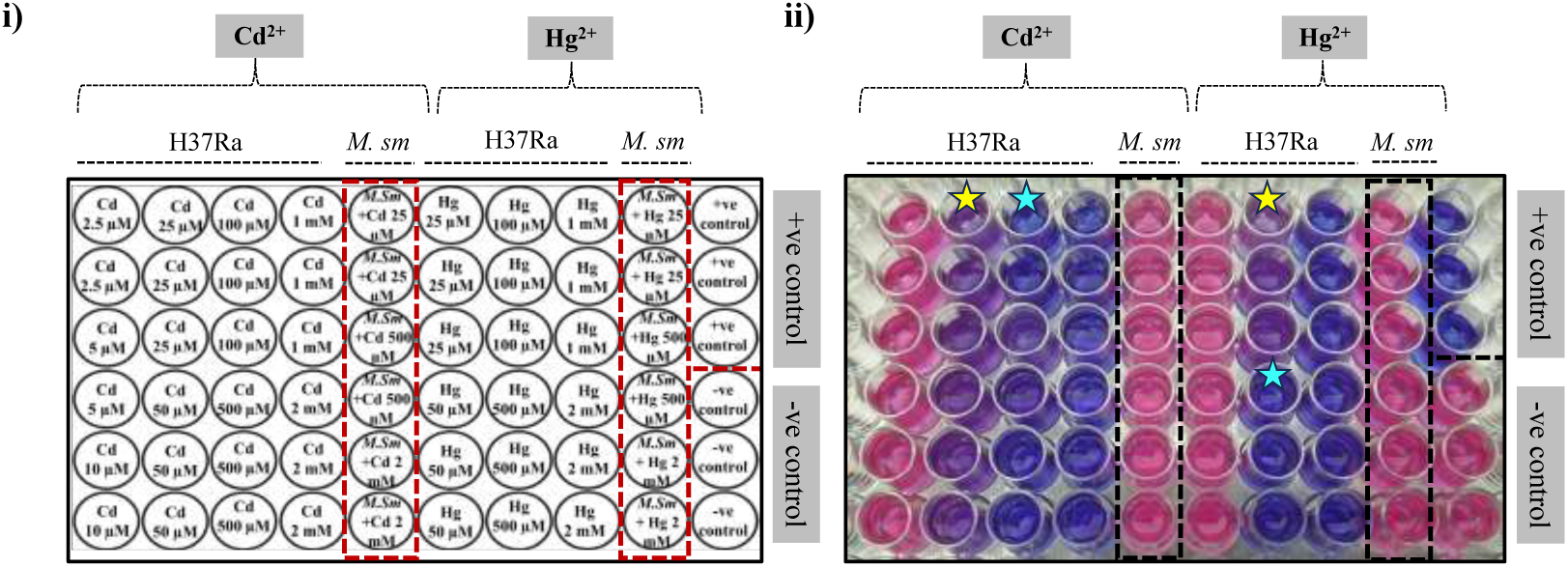
96-well plate-based biosensor assay demonstrating whole cell mycobacteria (*Mtb* H37Ra strain) as the biological indicator for toxic metal ions (Cd^2₊^ and Hg^2₊^) (i) Schematic representation of 96 well plate-based Biosensor platform. (ii) H37Ra cells (10^8^ CFU/ml) were seeded in the presence of varied concentration ranges of CdCl_2_ (2.5 µM- 2 mM) and HgCl_2_ (25 µM- 2 mM), respectively. Cells incubated in the presence of CdCl_2_ (2.5 µM- 10 µM) and HgCl_2_ (25 µM- 50 µM) were pink in color indicating healthy viable cells resisting the splicing inhibitory effect of the metal ions. In contrast, cells exposed to CdCl_2_ (25 µM and 50 µM) and 100 µM of HgCl_2_ displayed purple color suggesting partial loss of cell viability. Cells incubated with 100 µM-2 mM of CdCl_2_ and 100 µM- 2 mM of HgCl_2_ remained blue, suggesting complete loss of viable cell population. For a baseline correction, test organisms were allowed to grow in the absence of toxic metal ions and denoted as live cell control (negative control). SufB intein-less organism *Mycobacterium smegmatis (M. sm)* was used as additional experimental control and treated under similar conditions. *M. sm* cells exhibited pink color across a varied range of toxic metal ions, suggesting their cell viability as an intein-independent phenomenon. The experiments were performed in triplicates to confirm the reproducibility of data. Cyan stars-complete loss of viability (blue) & Yellow stars-partial loss of viability (purple).

Next, cells incubated in the presence of 25 µM and 50 µM of Hg^2₊^ resulted in healthy viable (pink) cell population, whereas cells in the presence of 100 µM Hg^2₊^ exhibited purple color, highlighting partial loss of cell viability at MIC value. Over rest of the concentration range for Hg^2₊^ (500 µM-2 mM), non-viable (blue) cells were noticed. These results were supported by SufB splicing attenuation by Hg^2₊^ (500 µM-2 mM), while a lower concentration (2.5 µM-10 µM) Hg^2₊^ failed to exert any such effects (Fig. 1, Supplemental Fig. S4). Although 100 µM of Hg^2₊^ didn’t influence SufB splicing, the N-terminal cleavage reaction was reduced significantly, marking this as the MIC value with partial loss of cell viability. Summarizing, persistence of blue color in the presence of Cd^2₊^ (100 µM until 2 mM), and Hg^2₊^ (500 µM until 2 mM) may be correlated to complete loss of H37Ra viability due to inhibition of *Mtb* SufB splicing and N-terminal cleavage reactions (Fig. 1). These results suggest the potential use of mycobacterial (H37Ra) cells as the biological indicator for a qualitative toxic metal detection.

Next, a quantitative analysis for the toxic metal ions was performed by measuring the absorbance of Alamar Blue reagent at 570 nm with 600 nm as a reference wavelength (74). Metal samples of known concentrations were prepared and loaded on to the 96-well plate and treated with Alamar Blue reagent under optimised experimental conditions [Fig. 3 (i)-(iv)]. A standard graph was generated by plotting the absorbance values of Alamar reagent vs the known concentrations of test metal ions (Cd^2₊^ and Hg^2₊^) [Fig. 3 (v) and (vi)]. Subsequently, a linear regression analysis was performed to determine the equation of the line, expressed as y = mx ₊ c. To further validate the standard plot, unknown samples with varying concentrations of Cd^2₊^ ions and Hg^2₊^ ions were prepared (Experimental Procedure). Absorbance values of the unknown samples were extrapolated onto standard graph to determine the concentrations of Cd^2₊^ and Hg^2₊^ (Fig. 3 (v), (vi) and Table S1). Unknown samples with concentration higher than the standard range may be adjusted via dilution, or the standard calibration plot may be extended to higher concentration ranges (128). However, sample concentration at a value lower than the standard range may not be determined accurately possibly due to no effect on intein splicing and hence no loss of cell viability.

**Fig. 3.**
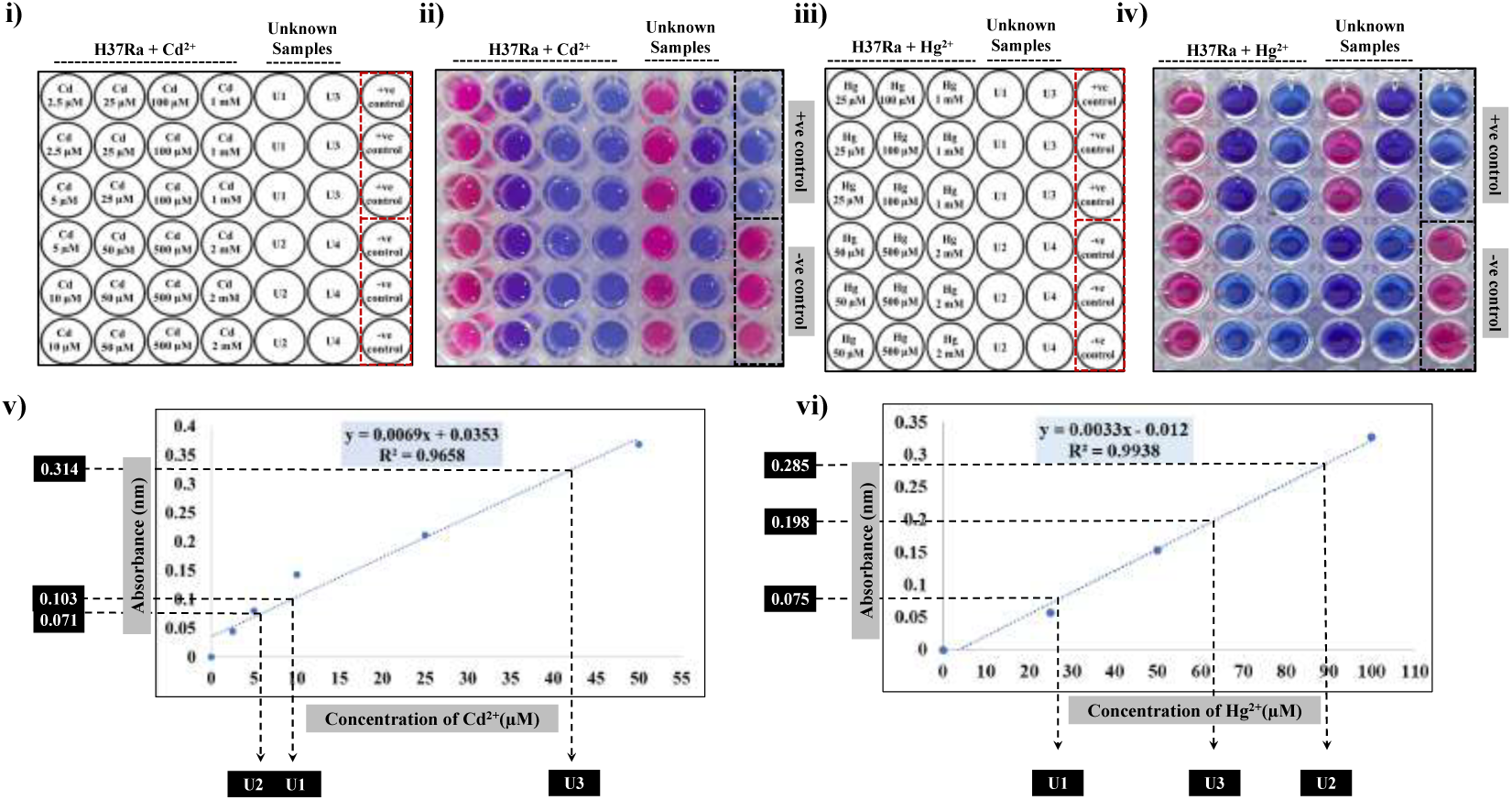
Standard curve for quantitative determination of Cd^2₊^ and Hg^2₊^ concentrations. (i), and (iii) Schematic representations of H37Ra-based Biosensor platform to detect CdCl_2_ and HgCl_2_. (ii), and (iv) *Mtb* H37Ra cells were grown in presence of varied concentration of CdCl_2_ and HgCl_2_. To establish a baseline, test organisms were allowed to grow in absence of toxic metal ions (live cell or negative control). (v), and (vi) The absorbance values of the Alamar Reagent solutions were plotted along the y-axis and the known concentrations of Cd^2₊^ and Hg^2₊^ were plotted along the x-axis. A linear regression analysis was conducted to obtain the equation for the line, and used to determine the concentration of unknown samples. The experiments were performed in triplicates to confirm the reproducibility of data.

Next, to elucidate the specificity of the designed biosensor platform, Cd^2₊^- and Hg^2₊^-specific metal chelators, Edetate Calcium Disodium (ECD), and meso-2,3-Dimercaptosuccinic acid (DMSA) were incubated alongside the test samples (129–134). Fig. 4 (i) is a schematic representation of the biosensor platform, where indicator mycobacterial cells were exposed to Cd^2₊^ and Hg^2₊^, in presence of Cd^2₊^-specific chelator, ECD. Fig. 4 (ii) demonstrates that ECD effectively resisted Cd^2₊^-induced loss of viability, resulting in healthy cells that transformed resazurin (blue) to resorufin (pink). Conversely, ECD failed to restore viability of Hg^2₊^ treated cells, where inhibition of SufB splicing, rendered non-viable (blue) cells. Next, mycobacterial cells were subjected to varying concentrations of Cd^2₊^ and Hg^2₊^, supplemented with metal chelator DMSA (Hg^2₊^ specific) [Fig. 4 (iii) and (iv)]. Hg^2₊^ treated H37Ra cells remained viable in presence of DMSA, while Cd^2₊^ continued its inhibitory effect on SufB splicing inducing non-viable (blue) cells. These observations affirmed the specificity of the designed biosensor platform to detect toxic metal ions Cd^2₊^ and Hg^2₊^.

**Fig. 4.**
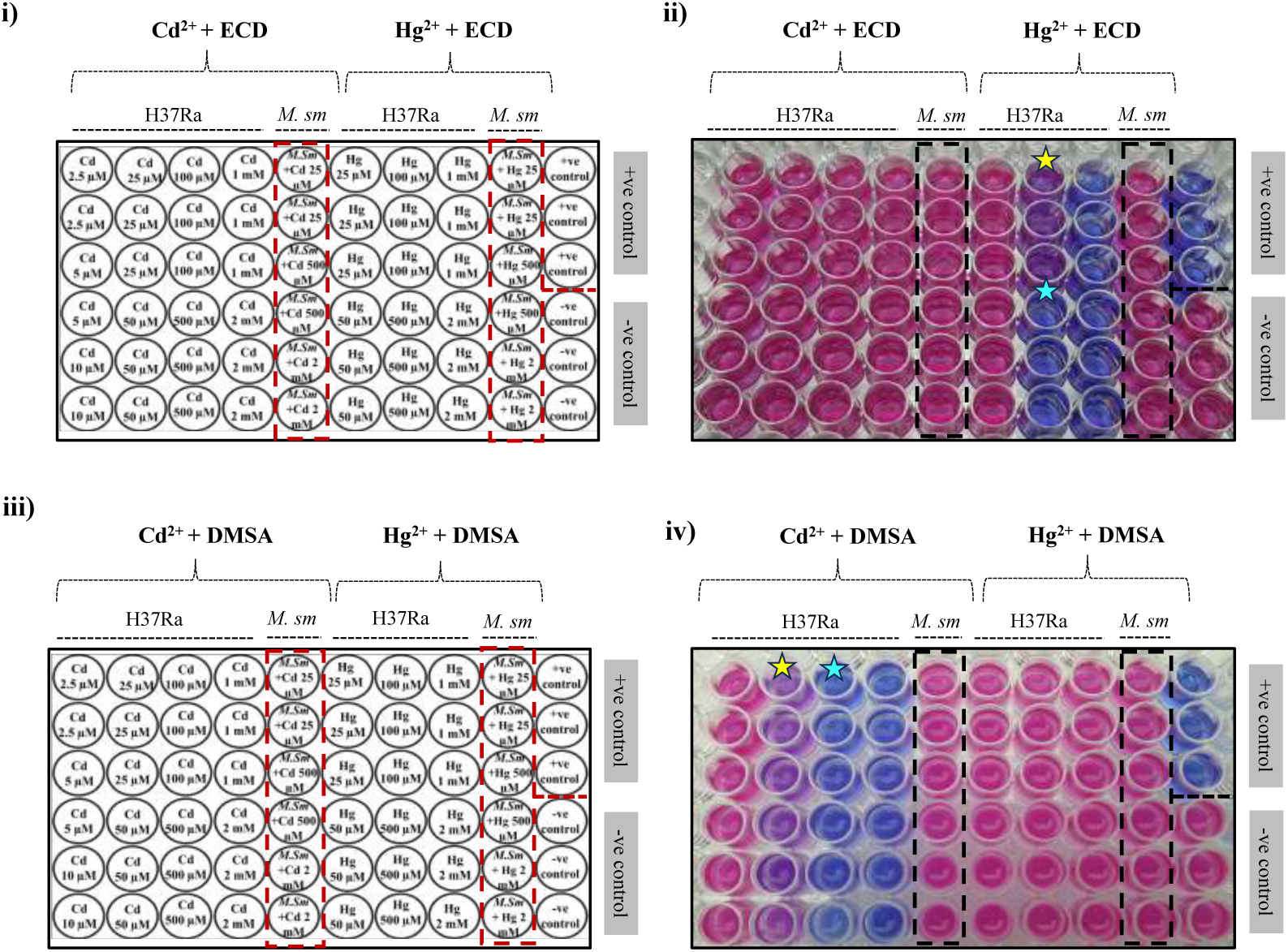
Determination of the specificity of *Mtb* H37Ra-based biosensor platform using metal chelators. (i) Schematic representation of the biosensor platform including Cd^2₊^ specific metal chelator (ECD) (ii) *Mtb* H37Ra cells (10^8^ CFU/ml) were allowed to grow in presence of varied concentration range of CdCl_2_ and HgCl_2_, along with ECD (1:1 ratio). ECD reversed the inhibitory effect of Cd^2₊^ inducing viable (pink) cells, whereas the effect of Hg^2₊^ persisted (iii) Schematic representation of the biosensor platform including Hg^2₊^ specific metal chelator (DMSA). (iv) *Mtb* H37Ra cells (10^8^ CFU/ml) were allowed to grow in the presence of varied concentration ranges of CdCl_2_ and HgCl_2_, including DMSA in the ratio 1:1. DMSA blocked the inhibitory effect of Hg^2^, rendering pink viable cells, whereas the effect of Cd^2₊^ persisted. For a baseline correction, H37Ra cells were grown in the absence of toxic metal ions as the live cell control. SufB intein-less organism *Mycobacterium smegmatis (M. sm)* was used as additional experimental control and treated under similar conditions. *M. sm* cells exhibited pink color across a varied range of toxic metal ions, suggesting their cell viability as an intein-independent phenomenon. The experiments were performed in triplicates to confirm the reproducibility of data. Cyan stars-complete loss of viability (blue) & Yellow stars-partial loss of viable cells (purple). ECD: Edetate Calcium disodium, DMSA: Meso-2,3-Dimercaptosuccinic acid.

Next for a multiplex metal screening process, the efficacy of the platform was assessed in presence of known splicing inhibitors Zn^2₊^ (100 µM-2 mM), Cu^2₊^ (100 µM-2 mM), Pt^4₊^ (20 µM-1 mM) along with Cd^2₊^ (10 µM-2 mM) and Hg^2₊^ (25 µM-2 mM) (Fig. 5) (43). Baseline correction was achieved by growing H37Ra cells in the absence of toxic metal ions (negative control), while SufB intein-less *M. sm* cells served as additional experimental control. Cells treated with 100 µM ZnCl_2_ and CuCl_2_, 20 µM PtCl_4_, 10 µM CdCl_2,_ and 25 µM HgCl_2_, turned pink in color, indicating intact viability. At intermediate concentrations, 600 µM of ZnCl_2_ and CuCl_2_, 40 µM PtCl_4_, 25 µM CdCl_2,_ and 100 µM HgCl_2_, a transition from blue to purple color was observed, suggesting partial loss of cell viability. Complete loss of viability was noted at higher concentration range, for ZnCl_2_ and CuCl_2_ (800 µM- 2 mM), PtCl_4_ (80 µM- 1 mM), CdCl_2_ (100 µM- 2 mM), and HgCl_2_ (500 µM- 2 mM) (Fig. 5 ii). Dead non-viable cells induced by Zn^2₊^, Cu^2₊^, and Pt⁴₊ may be correlated to inhibition of *Mtb* SufB splicing and N-terminal cleavage reactions as published earlier (43). In contrast, SufB intein-less organism (*M. sm*) remained viable (pink) under similar experimental conditions, suggesting cell viability as a splicing independent phenomenon. These observations may also be used to rule out metal toxicity on cell viability over the test concentration range.

**Fig. 5.**
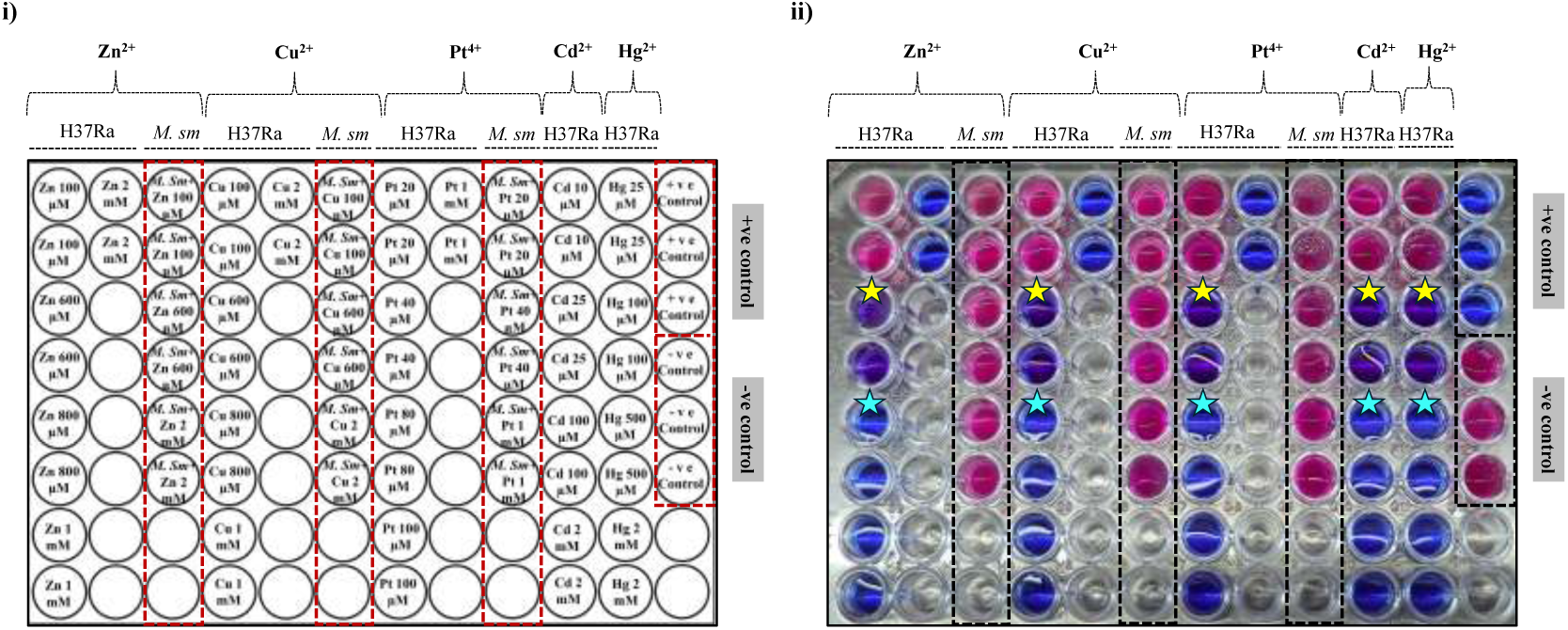
Evaluation of *Mtb* H37Ra-based biosensor platform for high-throughput and multiplex detection of trace metal elements and toxic metal ions (Zn^2₊^, Cu^2₊^, Pt^4₊^, Cd^2₊^, and Hg^2₊^). (i) Schematic representation of the multiplexing biosensor platform. (ii) H37Ra cells (10^8^ CFU/ml) were allowed to grow in the presence of ZnCl_2_ (100 µM- 2 mM), CuCl_2_ (100 µM- 2 mM), PtCl_4_ (20 µM- 1 mM), CdCl_2_ (10 µM- 2 mM) and HgCl_2_ (25 µM- 2 mM), respectively. Cells incubated with 100 µM of ZnCl_2_ and CuCl_2_, 20 µM of PtCl_4_, 10 µM of CdCl_2_ and 25 µM of HgCl_2_ were pink in color indicating healthy viable cells. In contrast, cells exposed to 600 µM of ZnCl_2_ and CuCl_2_, 40 µM of PtCl_4_, 25 µM of CdCl_2_ and 100 µM of HgCl_2_ turned purple in color suggesting partial loss of cell viability, whereas cells exposed to 600 µM- 2 mM of ZnCl_2_ and CuCl_2_, 80 µM- 1 mM of PtCl_4_, 100 µM- 2 mM of CdCl_2_ and 500 µM- 2 mM of HgCl_2_ remained blue in color suggesting complete loss of viable cell population. For a baseline correction, H37Ra cells were grown in the absence of toxic metal ions as the live cell control (negative control). SufB intein-less organism *Mycobacterium smegmatis (M. sm)* was used as additional experimental control and treated under similar conditions. *M. sm* cells exhibited pink color across a varied range of toxic metal ions, suggesting their cell viability as an intein-independent phenomenon. The experiments were performed in triplicates to confirm the reproducibility of data. Cyan stars-complete loss of viability (blue) & Yellow stars-partial loss of viability (purple).

The multiplexing ability of the biosensor platform was further examined by employing metal-specific chelators along with the metal-treated samples. The Cd^2₊^-specific chelator ECD and Hg^2₊^-specific chelator DMSA were tested alongside the broad-spectrum metal chelator EDTA. Fig. 6 (i) is a schematic representation of the biosensor platform, where indicator mycobacterial cells were incubated with Cd^2₊^-specific chelator, ECD. Fig. 6 (ii) demonstrates that co-incubation with ECD effectively reversed Cd^2₊^-induced loss of viability, restoring pink color. This effect was not observed in cells exposed to other metal ions, where ECD could not block metal induced inhibition. Similarly, DMSA reversed the toxic effects of Hg^2₊^ [Fig. 6 (iii) and (iv)], while other metal ions continued to attenuate cell viability. Notably, Since EDTA is a broad-spectrum metal chelator, it restored viability across all metal-treated samples [Fig. 6 (v) and (vi)]. Collectively, these findings validate the high-throughput and multiplexing capability of the developed biosensor platform, underscoring its potential for simultaneous detection of multiple toxic metal ions with high specificity and sensitivity.

**Fig. 6.**
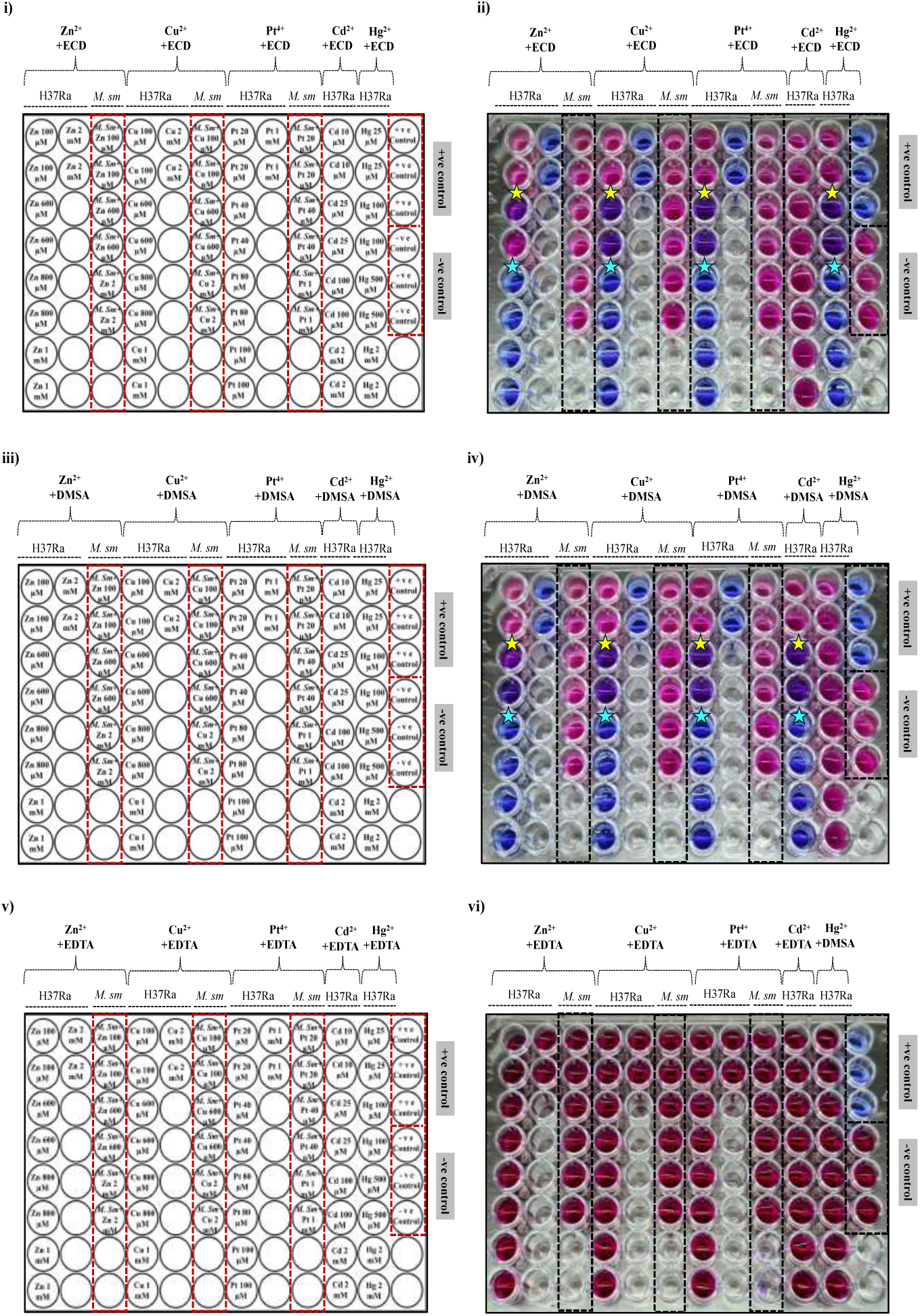
Determination of specificity of the *Mtb* H37Ra-based multiplexing biosensor platform. (i) Schematic representation of the multiplexing biosensor platform along with Cd^2₊^ specific metal chelator (ECD) (ii) *Mtb* H37Ra cells (10^8^ CFU/ml) were incubated with different concentration of ZnCl_2_, CuCl_2_, PtCl_4_, CdCl_2_ and HgCl_2_, along with ECD (1:1 ratio). ECD reversed the inhibitory effect of Cd^2₊^ rendering viable pink cells, whereas the effects of other metal ions persisted (iii) Schematic representation of the multiplexing biosensor platform along with Hg^2₊^ specific metal chelator (DMSA). (iv) *Mtb* H37Ra cells (10^8^ CFU/ml) were incubated with different concentration of ZnCl_2_, CuCl_2_, PtCl_4_, CdCl_2_ and HgCl_2_, along with DMSA in the ratio 1:1. DMSA inhibited the effect of Hg^2₊^ inducing viable pink cells, whereas the effect of other metal ions persisted. (v) Schematic representation of plate-based biosensor platform in presence of common metal chelator EDTA. (vi) *Mtb* H37Ra cells (10^8^ CFU/ml) were allowed to grow in presence of ZnCl_2_, CuCl_2_, PtCl_4_, CdCl_2_ and HgCl_2_ along with EDTA in the ratio 1:1. The inhibitory effects of test metal ions on mycobacterial cells were reversed by EDTA. For a baseline correction, H37Ra cells were grown in the absence of toxic metal ions, as the live cell control. SufB intein-less organism *Mycobacterium smegmatis (M. sm)* was used as additional experimental control and treated under similar conditions. *M. sm* cells exhibited pink color across a varied range of toxic metal ions, suggesting their cell viability as an intein-independent phenomenon. The experiments were performed in triplicates to confirm the reproducibility of data. Cyan stars-complete loss of viability (blue) & Yellow stars-partial loss of viable cells (purple). ECD: Edetate Calcium disodium, DMSA: Meso-2,3-Dimercaptosuccinic acid, EDTA: Ethylenediaminetetraacetic Acid.

In toto, the designed biosensor has demonstrated the potential of whole-cell *Mtb* H37Ra as an indicator to detect toxic metal ions as well as TMEs in test samples. The specificity of the designed metal-sensor was shown by the reversal of the inhibitory effects in the presence of Cd^2₊^- and Hg^2₊^-specific metal chelators, ECD and DMSA, respectively. Detection of metal ions at a concentration of 25 µM (Cd^2₊^) highlights the sensitivity of the biosensor platform, although it requires further optimization to cover the lower concentration range of various metal ions. We propose the use of whole-cell microorganisms carrying metal-sensing intein-bearing proteins as an indicator for the presence of toxic metal ions in environmental samples. This is a user-friendly method that can be performed in a 96-well microtiter plate, which would provide a color change between blue and pink while sensing metal ions. Since the activity of metal-sensing intein-carrying protein sequences are unique for the indicator (whole-cell) organisms, in the unlikely event of contamination in a biological facility, it should not raise concern about the validity of the results.

## Conclusion

Heavy metal ions are persistent environmental contaminants that pose significant risks to both ecological systems and human health. With the rising burden of metal pollution across diverse environmental and biological matrices, there is a pressing need for sensitive, specific, and user-friendly biosensing technology. Intein-mediated biosensors are emerging as promising tools in this context, particularly due to their unique splicing mechanism that operates efficiently under physiological conditions, eliminating the need for external cofactors or energy sources (34, 135). In this study, an innovative biosensor platform was developed, employing native H37Ra cells where intein splicing inhibition is a readout mechanism for the detection of toxic heavy metal ions (e.g., Cd^2₊^ and Hg^2₊^) and other trace metal elements (Fig. 7). The resulting alteration in mycobacterial viability is captured through a rapid and quantifiable resazurin-based colorimetric assay that reports color change between blue-purple-pink. Importantly, the platform is adaptable for multiplex detection of metal ions, including Zn^2₊^, Pt^4₊^, Cu^2₊^, Cd^2₊^, and Hg^2₊^, thereby offering a high-throughput solution for complex sample screening (Fig. 5 and 6). Further, this can be extended to sense metal ions by other whole-cell microorganisms, harbouring unique metal-sensing intein-containing proteins, hence offering a qualitative as well as quantitative detection method.

**Fig. 7.**
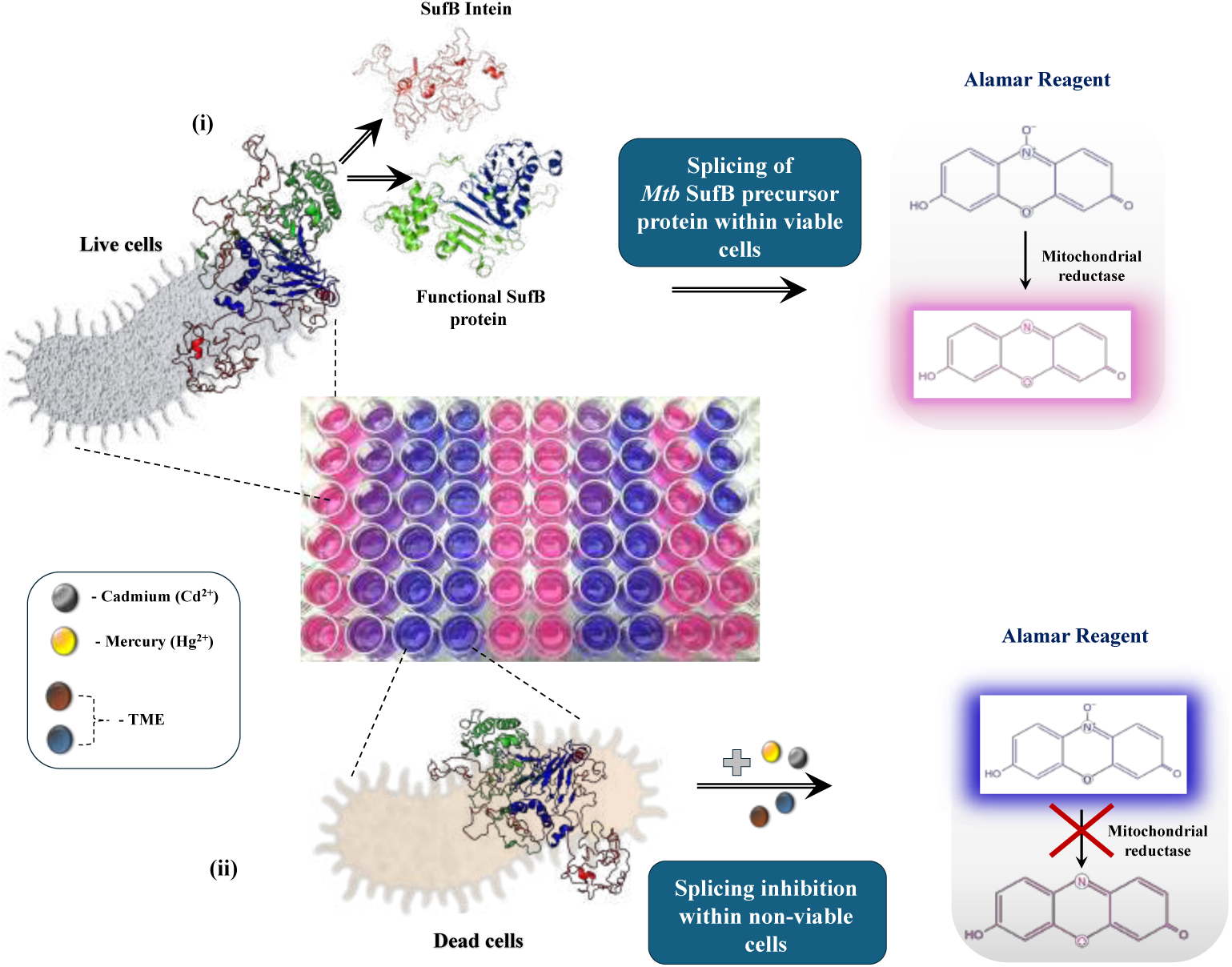
Schematic representation explaining the principle of *Mtb* H37Ra-based multiplexing biosensor platform. Whole-cell mycobacteria (H37Ra) were used as biological indicator for the detection of toxic metal ions (Cd^2₊^ and Hg^2₊^) and other trace metal elements (TME). (i) Intein splicing under normal physiological conditions generates functional SufB protein, while viable mycobacterial cells convert resazurin (blue) to resorufin (pink). (ii) In the presence of toxic metal ions Cd^2₊^ and Hg^2₊^ and TME, SufB intein splicing is inhibited resulting in non-viable mycobacterial cells, thus there is no conversion of resazurin (blue) to resorufin (pink).

SufB is a critical component of SUF system, the exclusive pathway for Fe-S cluster assembly in mycobacteria. Previous studies have also highlighted the pivotal role of *Mtb* SufB protein in Fe-S cluster biogenesis, and mycobacterial survival (61, 136). Since the functionality of SUF protein complex is dependent on generation of active SufB protein via splicing, splicing inhibition presents a promising approach for antimicrobial intervention by targeting mycobacterial viability. Several studies have shown that divalent cations (Zn^2₊^ and Cu^2₊^), ROS, RNS, chlorine derivatives, small molecules (cisplatin and diethanolamine) and zinc oxide nanoparticles can act as potent splicing inhibitors. These agents have influenced mycobacterial survival, further validating the relevance of targeting intein splicing for antimycobacterial drug development (40, 43, 51, 56, 58, 64, 66, 137–139). Earlier work from our group has demonstrated that metal ions such as Cu^2₊^, Zn^2₊^, and Pt⁴₊ inhibit SufB splicing, establishing a correlation between metal-induced stress and altered intein splicing activity in *Mtb* (43). Building on these findings, the current work explores regulatory roles of toxic metal ions on *Mtb* SufB splicing and cleavage activity. Alamar Blue assay was employed to examine the influence of TMEs and toxic metal ions on the viability of *Mtb* cells. The outcome of this research will advance the real time application of H37Ra/*Mtb* based whole-cell biosensor to detect wide range of metal ions. The platform described herein is robust, enabling multiplex screening of metal ions while maintaining specificity and sensitivity. Importantly, cellular viability loss in response to metal exposure is linked to an observable color shift between blue and pink, providing a quantifiable readout for metal ion detection in test samples.

In this study, splicing inhibition was evaluated both *in vitro*, using purified SufB protein and *in vivo* within H37Ra cells in a whole-cell biosensor platform, where the observed effects on mycobacterial viability were correlated to SufB splicing inhibition. Given that H37Ra is a slow-growing strain with a mean generation time of approximately 44 hours, the experimental window was established at four days to encompass at least two full generations, permitting effective assessment of metal-induced viability changes (140–142). Notably, the logarithmic growth phase of mycobacteria extends from four to twenty days, ensuring that the platform remains adaptable to varied experimental requirements. To further strengthen our hypothesis, SufB intein less organism, *M. sm* control was included; viability of *M. sm* persisted throughout the experimental period, suggesting that the tested metal ions do not elicit growth inhibitory or toxic effects in the absence of SufB intein. *Mtb* contains three intein containing proteins; SufB, RecA and DnaB. While SufB is essential for Fe-S cluster assembly, RecA and DnaB play pivotal roles during DNA replication; taken together these proteins are critical for mycobacterial survival (143–145). Although current study explains loss of mycobacterial cell viability as a result of inhibition of *Mtb* SufB splicing, the possibility of RecA and DnaB splicing inhibition by the test metal ions cannot be ruled completely. Currently, there are no literatures to show inhibitory effect of metal ions other than Zn^2₊^ on *Mtb* DnaB splicing (49). Moreover there are alternative helicases present in mycobacteria to take over the function of DnaB (123–125). Several studies have reported regulatory roles of Zn^2₊^, Cd^2₊^, Co^2₊^, Ni^2₊^, Cu^₊^/Cu^2₊^, CisPt ions on *Mtb* RecA intein splicing (45, 47, 58, 137). However, current study has not evaluated the RecA intein inhibition in mycobacteria. Hence, this approach offers a multiplexing biosensor unit which can accommodate cell viability detection via splicing inhibition in many organisms carrying different metal-sensitive intein containing proteins simultaneously.

The use of non-genetically modified *Mtb* H37Ra cells ensures biosecurity compliance while providing a biologically relevant sensing chassis. These mycobacterial cells are not only easy to cultivate and maintain but also exhibit remarkable long-term viability, supporting extended shelf-life and storage without specialized conditions. The platform also demonstrates good sensitivity in detecting metal ions over a broad dynamic range, and substrate specificity, making it suitable for metal ion screening in standard laboratory setups.

However, some limitations persist such as complex environmental or biological samples containing mixtures of multiple metal ions may result in signal overlap or cross-reactivity with variable binding affinities. Furthermore, while microbial whole-cell biosensors offer an excellent alternative for initial screening and field deployment, advanced physicochemical instruments remain crucial when trace-level quantification is required, where intein activity may not be efficiently regulated.

In summary, our findings highlight the potential of whole cell intein-based biosensors as user-friendly, and cost-effective tools for the detection of toxic metal and trace metal ions. Future work should aim to refine their activity, enhancing environmental robustness, and advancing the platform from lab-scale to deployable technologies. This can be accelerated through collaborative efforts across academic, industrial, and governmental sectors.

## Abbreviation

Ni-NTA: Nickel-nitrilotriacetic acid
TCEP.HCl: Tris 2-carboxyl ethyl phosphine. Hydrochloric acid
IPTG: Isopropyl β-d-1-thiogalactopyranoside
Trp: Tryptophan
SDS: Sodium Dodecyl Sulfate
PAGE: Polyacrylamide gel electrophoresis
ICP-OES: Inductively Coupled Plasma Optical Emission Spectroscopy
UV-Vis: UV-visible spectroscopy
PVDF: Polyvinylidene fluoride
BSA: Bovine serum albumin
*Mtb*: *Mycobacterium tuberculosis*
WT: Wild type
SI: Splicing inactive
Cd^2₊^: Cadmium
Hg^2₊^: Mercury
Pb^2₊^: Lead
Cr^3₊^: Chromium
Cu^2₊^: Copper
Zn^2₊^: Zinc
Pt^4₊^: Platinum

## Data availability

The data underlying the present article are available in the article and in its online Supplementary materials. Additional data will be made available on request if needed.

## Supplemental Data

This article contains supplemental data.

## Acknowledgment

The plasmids used in this study are borrowed from Prof. Marlene Belfort’s Lab, SUNY, Albany, NY, USA. Our sincere gratitude to the Central Instrumentation Facility (CIF) at OUAT (Odisha University of Agriculture & Technology), Bhubaneswar, and Dr. S.K. Dash for his help in conducting the ICP-OES analysis at OUAT, Bhubaneswar.

## Author Contributions

**AM:** Writing – Original Draft, Writing - Review & Editing, Methodology, Investigation, Formal analysis, Data curation, Visualization, Validation, Software, Resources, Funding acquisition; **AN:** Conceptualization, Methodology, Investigation, Formal analysis, Data curation; Writing - Review & Editing,; **SN:** Conceptualization, Writing - Original Draft, Writing - Review & Editing, Project administration, Supervision, Methodology, Investigation, Resources, Funding acquisition.

## Funding and Additional Information

Ms. Ashwaria Mehra is supported by INSPIRE fellowship (DST/INSPIRE/03/2021/002800/IF 190921); INSPIRE Division, DST, Government of India. This work was also supported by the UGC-DAE consortium for scientific research (UGC-DAE-CSR-KC/CRS/15/IOP/08/0562), Kolkata, India.

## Conflict of Interest

The authors declare that they have no known competing financial interests or personal relationships that could have appeared to influence the work reported in this paper.

## Notes

### Competing Interest Statement

The authors have declared no competing interest.

### Summary of Updates

This version of the manuscript has been revised to update the following: 1. Abstract has also been modified. 2. Introduction has been modified to address the novelty of the current study 3. We have also added research questions in the introduction

