## Supplemental Data for "Designing a robust whole-cell biosensor for detection of toxic metals using intein splicing inhibition of *Mycobacterium tuberculosis* SufB protein"

**Running Title:** Intein-based whole-cell biosensor detects toxic metals

| **Table of Contents** | **Page No.** |
|---|---|
| **Figure S1.** Schematic diagram illustrating different steps of *Mtb* SufB precursor protein splicing, and cleavage reactions. | S3 |
| **Figure S2.** (A) Western Blot analysis confirming the minimum effective concentrations of toxic metal ions on *Mtb* SufB precursor protein splicing and N-terminal cleavage reactions. | S4-S5 |
| **S2.** (B) Evaluation of toxic metal ion effect on *Mtb* SufB precursor protein degradation. | S4-S5 |
| **Figure S3.** Gradient assay to evaluate the effect of lower concentrations of $CdCl_2$ (2.5 μM-100 μM) on *Mtb* SufB precursor protein splicing and N-terminal cleavage reactions. | S6 |
| **Figure S4.** Gradient assay to evaluate the effect of lower concentrations of $HgCl_2$ (2.5 μM –100 μM) on *Mtb* SufB precursor protein splicing and N-terminal cleavage reactions. | S7-S8 |
| **Figure S5.** Effect of $CdCl_2$ and $HgCl_2$ on *Mtb* SufB precursor protein splicing and N-terminal cleavage reactions over varying period. | S9 |
| **Figure S6.** UV-visible spectroscopic analysis to detect the interaction between toxic metal ions and *Mtb* SufB precursor protein. | S10 |
| **Figure S7.** Tryptophan fluorescence assay to evaluate the interaction between the *Mtb* SufB precursor protein and toxic metal ions. | S11 |
| Quantitative evaluation of *in vitro* biosensor platform for the detection of toxic metal ions. | S12-13 |
| **Table S1.** The absorbance value of unknown samples and their corresponding concentrations as obtained from the linear regression plot. | S13 |

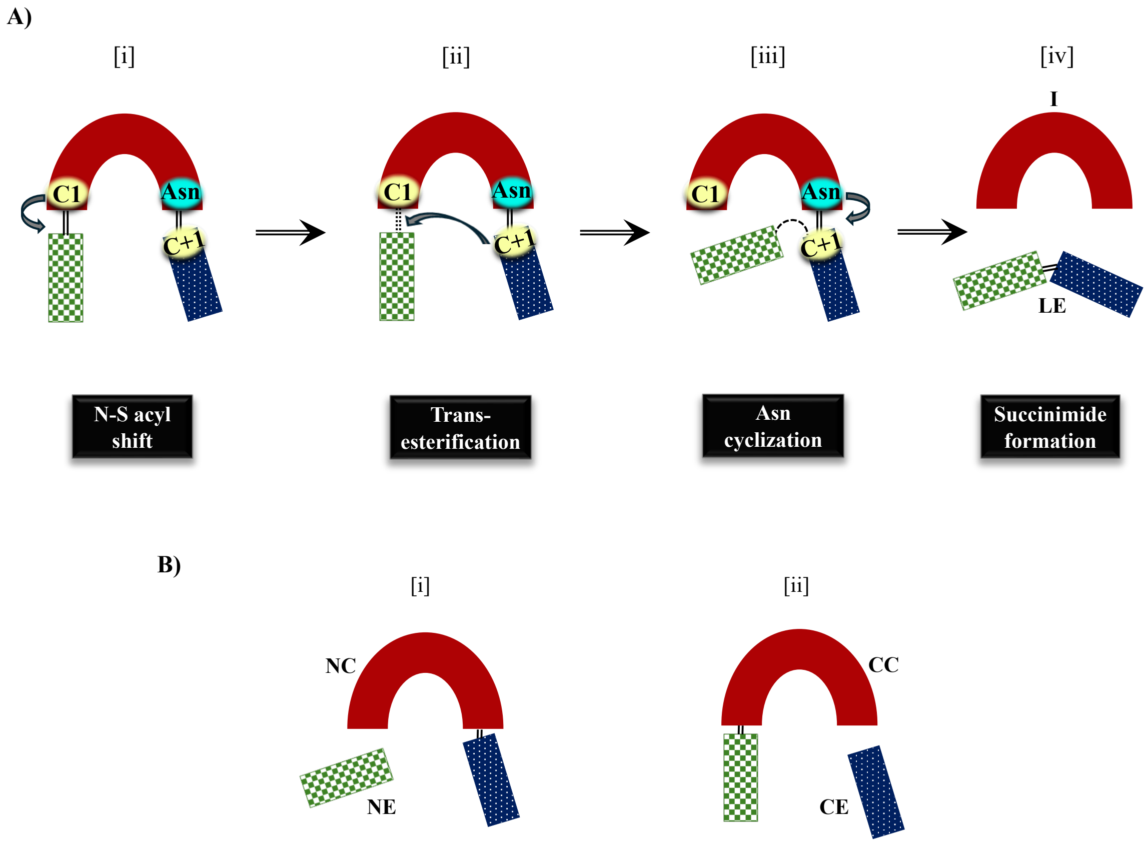

**Figure S1. Schematic diagram illustrating different steps of *Mtb* SufB precursor protein splicing and cleavage reactions. (A)** Canonical intein splicing proceeds through four sequential nucleophilic displacement reactions. (i) The first nucleophile, a C1 from A-block initiates the N-S acyl rearrangement at the N-terminal splice junction forming a thioester intermediate, (ii) Next, a nucleophilic attack by the C+1 residue of the c-extein drives transesterification, resulting in a branched intermediate, (iii) This branch intermediate is resolved by cyclization of the terminal Asn residue, promoting cleavage at the C-terminal splice site and (iv) Finally a second N-S acyl shift occurs leading to ligation of the flanking extein regions by an amide bond. **(B)** Off-pathway cleavage reaction product: (i) Cleavage at the N-terminal splice site leads to the release of the N-extein and an N-terminal cleavage product, (ii) Cleavage at the C-terminal site results in release of the C-extein and a C-terminal cleavage product. C1: cysteine1, Asn: Aspergine359, C+1: cysteine+1, I: Intein, LE: Ligated exteins, NC: N-terminal cleavage product, NE: N-extein, CC: C-terminal cleavage product, CE: C-extein.

**Western blot analysis to confirm the effect of toxic metal ions on *Mtb* SufB precursor protein splicing and cleavage reactions**

*In vitro,* refolding of the denatured SufB precursor protein was conducted for 4 hours in the presence of varying concentrations of metal ions ($Cd^{2+}$, $Hg^{2+}$, $Cr^{3+}$, $Pb^{2+}$) (Fig. 1, main text). Experimental details and discussions are provided in the main text. Western blot analysis (Figure S2) confirmed the identity of the various splicing and N-terminal cleavage products resolved through SDS PAGE (Fig. 1, main text).

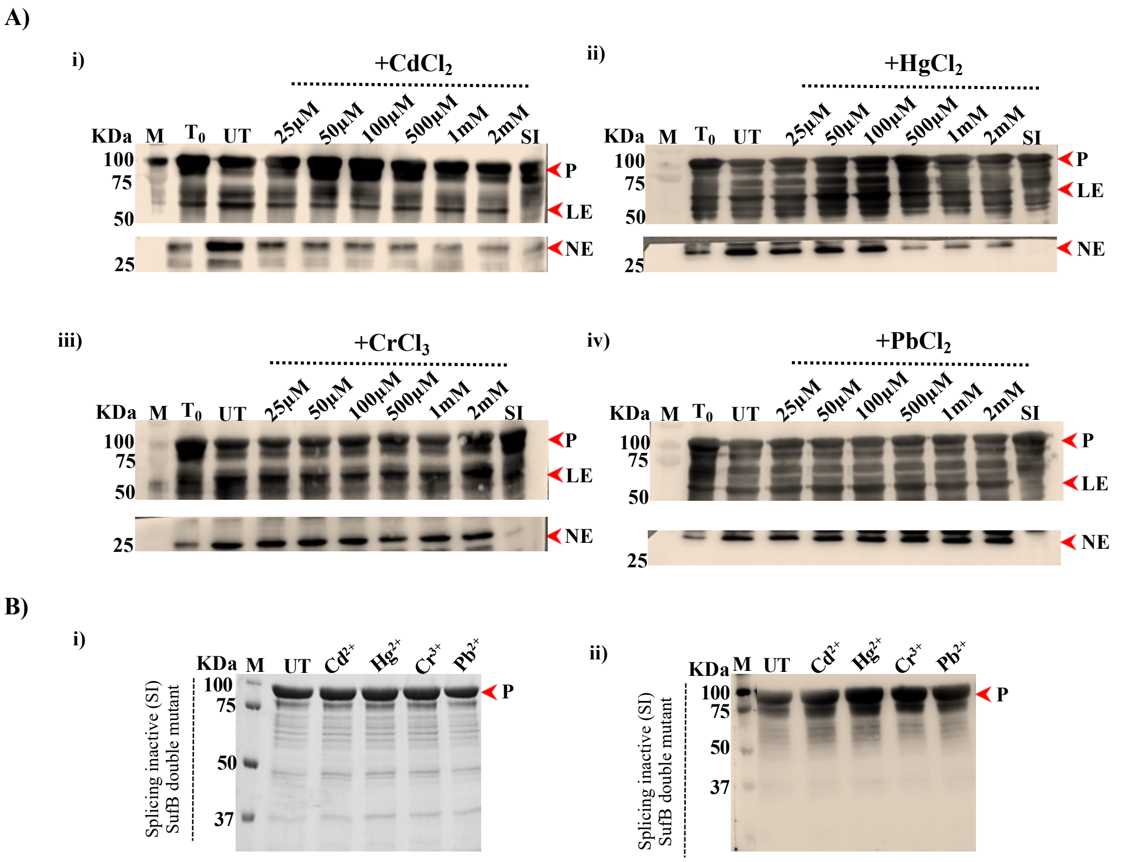

**Figure S2. (A) Western Blot analysis confirming the minimum effective concentrations of toxic metal ions on *Mtb* SufB precursor protein splicing and N-terminal cleavage reaction.** Products from the *in vitro* refolding assay in the presence of (i) $CdCl_2$, (ii) $HgCl_2$, (iii) $CrCl_3$, and (iv) $PbCl_2$ were resolved using 4–10% gradient SDS-PAGE, and the identity of the protein products (from Fig.1A, main text) were confirmed via western blot as shown here. Anti-6X(His) antibodies detected the presence of 6X(His)-tagged P, LE, and NE. NE is blotted separately using a higher concentration of the primary antibody. Lane 1 (M): Prestained protein molecular weight ladder; Lane 2: ($T_0$) splicing and N-terminal cleavage reactions at time 0; Lane 3 (UT): untreated sample, renatured without metal treatment, Lanes 4-9: *Mtb* SufB precursor refolded with varied concentrations of test metal ions. Lane 10: splicing inactive SufB double mutant (SI, C1A/N359A). P: Precursor; LE: Ligated extein; NE: N-extein. **(B) Evaluation of toxic metal ion effect on *Mtb* SufB precursor protein degradation.** (i) Splicing inactive (SI) SufB double mutant (C1A/N359A) was renatured in presence of toxic metal ions ($CdCl_2$, $HgCl_2$, $CrCl_3$, and $PbCl_2$) and resolved through SDS-PAGE. Un-spliced and intact SufB (C1A/N359A) precursor protein was observed indicating lack of protein degradation in presence of test metal ions. Lane 1 (M): Prestained protein molecular weight ladder; Lane 2 (UT): untreated sample; Lanes 3-6: Splicing inactive (SI) SufB double mutant (C1A/N359A) renatured in presence of toxic metal ions ($CdCl_2$, $HgCl_2$, $CrCl_3$, and $PbCl_2$). (ii) Western blot confirms the identity of protein products from Figure S2B(i). Anti-6X(His) antibody detected, 6X(His)-tagged un-spliced and intact SufB (C1A/N359A) precursor protein. Lane 1 (M): Prestained protein molecular weight ladder; Lane 2 (UT): untreated sample; Lanes 3-6: Splicing inactive (SI) SufB double mutant (C1A/N359A) renatured in presence of toxic metal ions ($CdCl_2$, $HgCl_2$, $CrCl_3$, and $PbCl_2$). P: *Mtb* SufB precursor double mutant (C1A/N359A).

**Gradient assay to evaluate the sensitivity of *Mtb* SufB precursor protein to lower concentration range of $CdCl_2$ (2.5 μM to 100 μM)**

To determine the effects of lower concentrations of $CdCl_2$ (2.5 μM –100 μM) on the N-terminal cleavage and splicing reactions of the *Mtb* SufB precursor protein, an *in vitro* gradient assay was conducted for 4 hours at 25 °C. *Mtb* SufB precursor protein was refolded in the presence of various concentrations of $CdCl_2$ (2.5 μM –100 μM) and the reaction products were resolved using 4–10% SDS-PAGE. Experimental details are provided in the main text (Materials and Methods). No significant effect was observed at 2.5 μM and 5 μM $CdCl_2$. However, 10 μM $CdCl_2$ induced mild but statistically significant reductions in % of splicing ($p = 0.02$) and % of N-terminal cleavage ($p = 0.002$) reactions. Significant splicing and cleavage inhibition was induced by $CdCl_2$ at higher concentrations (25, 50, and 100 μM). These results are discussed in the main text (Toxic metal ions inhibit *Mtb* SufB precursor protein splicing; Results and Discussion).

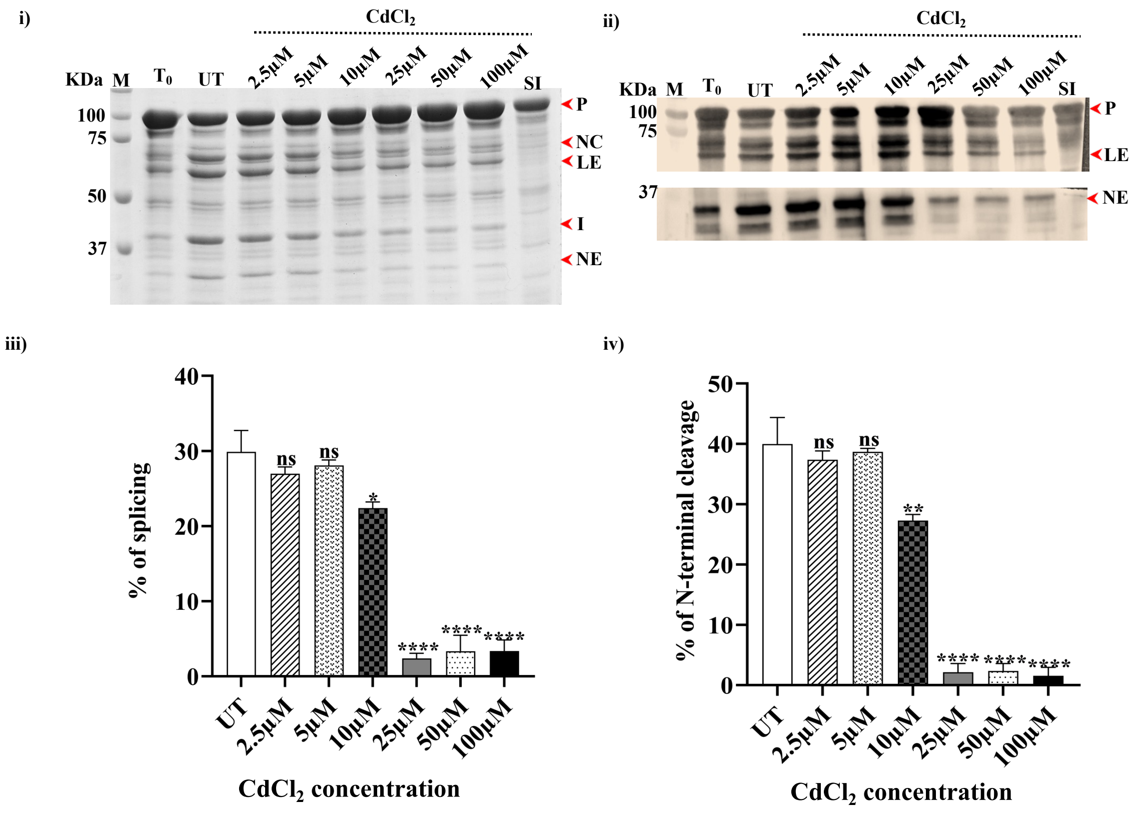

**Figure S3. Gradient assay to evaluate the effect of lower concentrations of $CdCl_2$ (2.5 µM –100 µM) on *Mtb* SufB precursor protein splicing and N-terminal cleavage reactions.** (i) Splicing and N-terminal cleavage reaction products following 4 hours of protein refolding were resolved using 4-10% gradient SDS-PAGE. Lane 1: M, Prestained protein molecular weight ladder; Lane 2: ($T_0$) splicing and N-terminal cleavage reaction at time 0; Lane 3: (UT) N-terminal cleavage and splicing reaction in untreated protein sample; Lanes 4-9: *Mtb* SufB precursor protein splicing and N-terminal cleavage products in presence of $CdCl_2$ (2.5µM- 100µM); Lane 10: (SI) Splicing-inactive SufB double mutant (C1A/N359A) as a negative control. (ii) Western blot confirms the identity of protein products from Figure S3(i). Anti-6X(His) antibodies detected the presence of 6X(His)-tagged P, LE, and NE. NE is blotted separately with a higher concentration of primary antibody and is shown separately. (iii) and (iv) Bar graphs depicting splicing and N-terminal cleavage efficiencies following $CdCl_2$ treatment. The data shown are extracted from Figure S3(i). These graphs were plotted after densitometric analyses using GelQuant.Net biochemical solutions. All the experiments were performed in triplicates, and error bars represent mean (±1) SEM from three independent sets of experiments. M: Prestained protein molecular weight ladder, P: Precursor, NC: N-terminal cleavage product, LE: Ligated extein, I: Intein, NE: N-extein.

**Gradient assay to evaluate the sensitivity of *Mtb* SufB precursor protein to lower concentration range of $HgCl_2$ (2.5 µM- 100 µM)**

To determine the effects of lower concentrations of $HgCl_2$ (2.5 µM –100 µM) on the N-terminal cleavage and splicing reactions of the *Mtb* SufB precursor protein, an *in vitro* gradient assay was conducted for 4 hours at 25 °C. *Mtb* SufB precursor protein was refolded in the presence of various concentrations of $HgCl_2$ (2.5 µM –100 µM), and the reaction products were resolved using 4–10% SDS-PAGE. Experimental details are provided in the main text (Materials and Methods). No significant effect was observed from 2.5 µM –100 µM of $HgCl_2$. Significant splicing and cleavage inhibition was induced by $HgCl_2$ at higher concentrations (500 µM, 1 mM, and 2 mM). These results are discussed in the main text (Toxic metal ions inhibit *Mtb* SufB precursor protein splicing and N-terminal cleavage reactions; Results and Discussion).

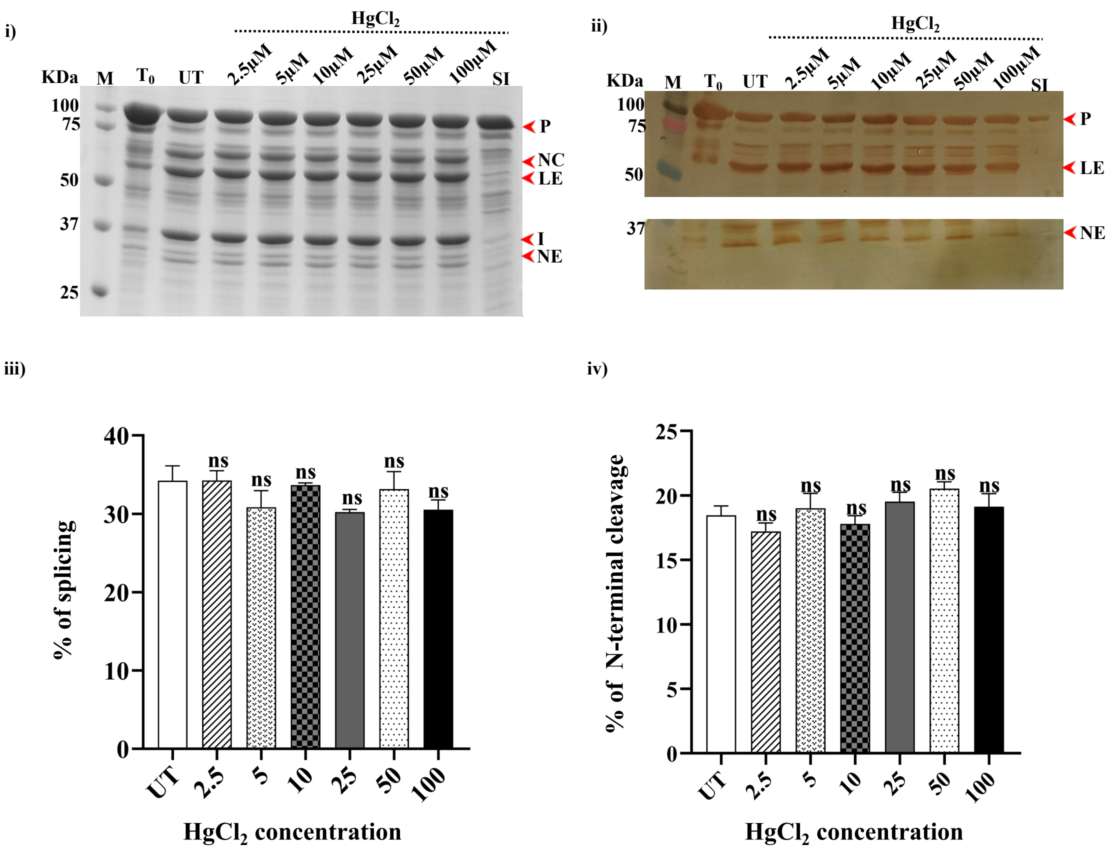

**Figure S4. Gradient assay to evaluate the effect of lower concentrations of $HgCl_2$ (2.5 µM –100 µM) on *Mtb* SufB precursor protein splicing and N-terminal cleavage reactions.** (i) Splicing and N-terminal cleavage reaction products following 4 hours of protein refolding were resolved using 4-10% gradient SDS-PAGE. Lane 1: M, Prestained protein molecular weight ladder; Lane 2: ($T_0$) splicing and N-terminal cleavage reaction at time 0; Lane 3: (UT) N-terminal cleavage and splicing reaction in untreated protein sample; Lanes 4-9: *Mtb* SufB splicing and N-terminal cleavage products in presence of $HgCl_2$ (2.5µM- 100µM); Lane 10: (SI) Splicing-inactive SufB double mutant (C1A/N359A) as a negative control. (ii) Western blot confirms the identity of protein products from Figure S4(i). Anti-6X(His) antibodies detected the presence of 6X(His)-tagged P, LE, and NE. NE is blotted separately with a higher concentration of primary antibody and is shown separately. Western blot images were developed using DAB substrate. (iii) and (iv) Bar graphs depicting splicing and N-terminal cleavage efficiencies following $HgCl_2$ treatment. The data shown are extracted from Figure S4(i). These graphs were plotted after densitometric analyses using GelQuant.Net biochemical solutions. All the experiments were performed in triplicates, and error bars represent mean (±1) SEM from three independent sets of experiments. M: Prestained protein molecular weight ladder, P: Precursor, NC: N-terminal cleavage product, LE: Ligated extein, I: Intein, NE: N-extein, DAB: 3,3′-diaminobenzidine.

**Effect of $CdCl_2$ and $HgCl_2$ on the splicing and N-terminal cleavage reactions of *Mtb* SufB precursor protein over varying period**

The effects of minimum effective concentration of $CdCl_2$ (25 µM ) and $HgCl_2$ (500 µM) on the splicing and N-terminal cleavage reactions of *Mtb* SufB protein were assessed over varying time (10 min to 4 hr.). *In vitro* refolding of denatured *Mtb* SufB precursor protein was performed in the presence and absence of the test metal ions over different period. Detailed experimental procedures and analyses are mentioned in the main text (Toxic metal ions inhibit *Mtb* SufB precursor protein splicing; Results and Discussion).

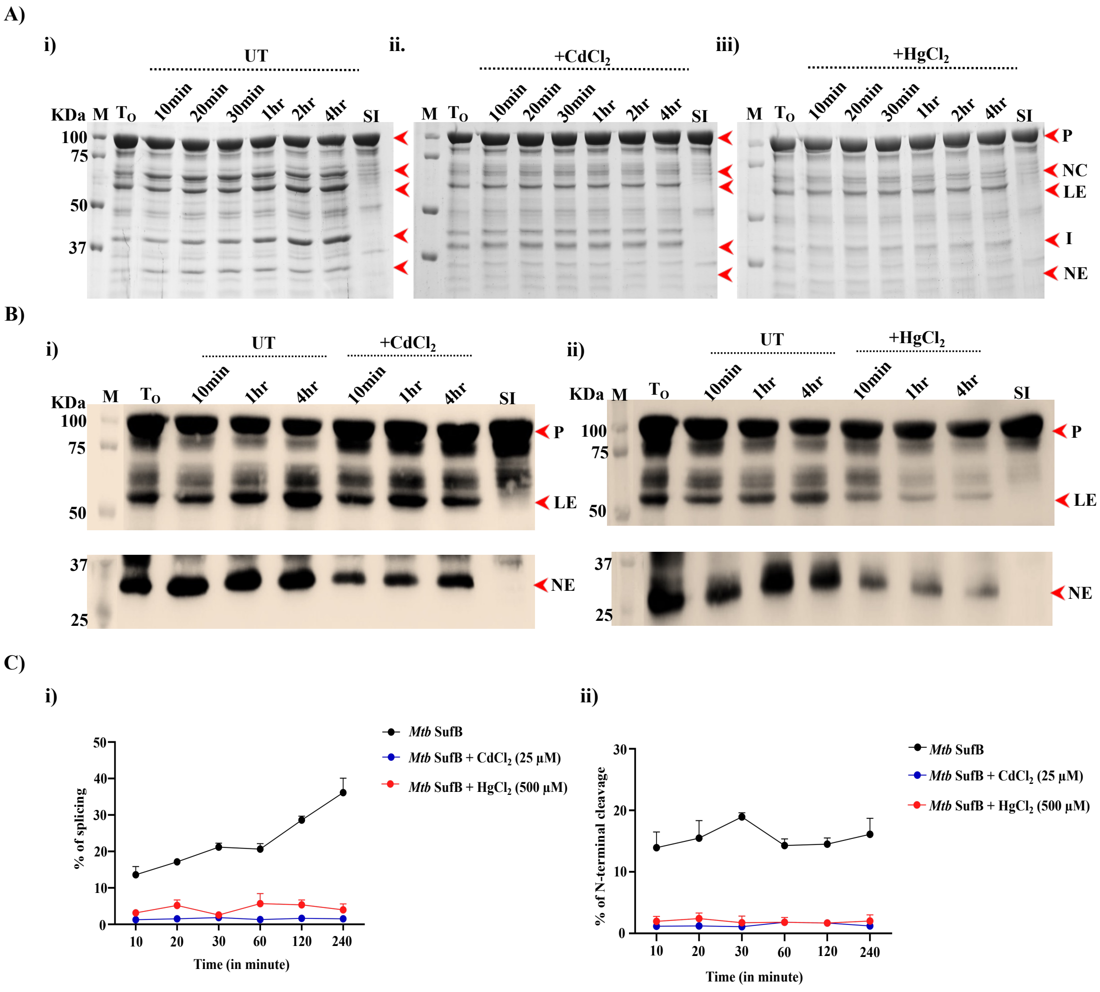

**Figure S5. Effect of $CdCl_2$ and $HgCl_2$ on *Mtb* SufB precursor protein splicing and N-terminal cleavage reactions over varying period.** (A) *In vitro* refolding of *Mtb* SufB precursor protein was conducted in the (i) absence of toxic metal (Untreated sample, UT) and the presence of (ii) $CdCl_2$ (25 µM) and (iii) $HgCl_2$ (500 µM) over varying period. N-terminal cleavage and splicing products were resolved through 4~10% gradient SDS-PAGE. (B) The identity of the protein products was confirmed via western blot using anti-6X(His) antibodies. The NE product was probed separately using a higher concentration of the primary antibody. P: Precursor; LE: Ligated extein; NE: N-extein. (C) Data are derived from Figure S5A, and line plots were plotted after densitometric analyses using GelQuant.Net biochemical solutions. Line plots presenting a comparative analysis of (i) % of splicing and (ii) % of N-terminal cleavage efficiencies in the presence and absence of $CdCl_2$ and $HgCl_2$ over different time. For baseline correction, values for 0 h were subtracted from each corresponding time point results. Graphs were generated using GraphPad Prism (version 5.01; GraphPad Software, San Diego, CA, USA; www.graphpad.com). All experiments were performed in triplicates; error bars represent the mean (± 1) SEM from three independent experiments. M: Prestained protein molecular weight marker; P: Precursor; NC: N-terminal cleavage product; LE: Ligated extein; I: Intein; NE: N-extein.

**Protein ~ metal interaction study**

<u>UV-Visible spectroscopic analysis</u>:

Protein~metal interactions were assessed by UV-visible spectroscopy in the presence and absence of toxic metal ions. Detailed experimental procedures and analyses are described in the main text (UV-Visible spectroscopic analysis; Results and Discussion).

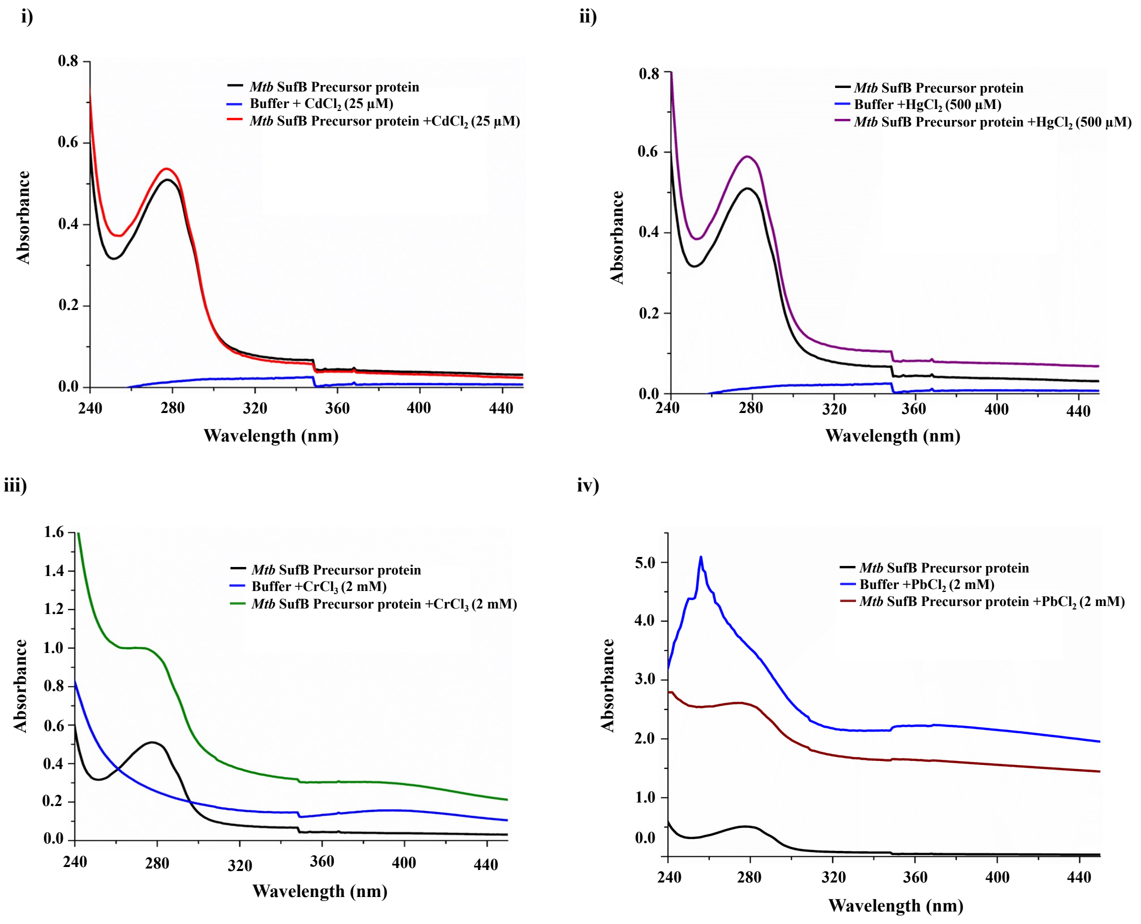

**Figure S6. UV-visible spectroscopic analysis to detect the interaction between toxic metal ions and *Mtb* SufB precursor protein.** Denatured *Mtb* SufB precursor protein was incubated with (i) $CdCl_2$, 25 µM (ii) $HgCl_2$, 500 µM (iii) $CrCl_3$, 2mM and (iv) $PbCl_2$, 2mM for 4 hours at 25ºC. Absorbance spectra were recorded over 200-800 nm. Changes in spectral profiles between untreated and metal-treated samples were analysed by plotting the absorbance values versus wavelength.

**Metal-induced alteration in tryptophan fluorescence (Tryptophan fluorescence assay):**

Tryptophan fluorescence spectroscopy was conducted to assess conformational changes in *Mtb* SufB precursor protein upon interaction with the toxic metal ions. Detailed experimental procedures and data analyses are provided in the main text (Tryptophan Fluorescence spectroscopy reveals metal-induced conformational changes in *Mtb* SufB precursor protein; Results and Discussion).

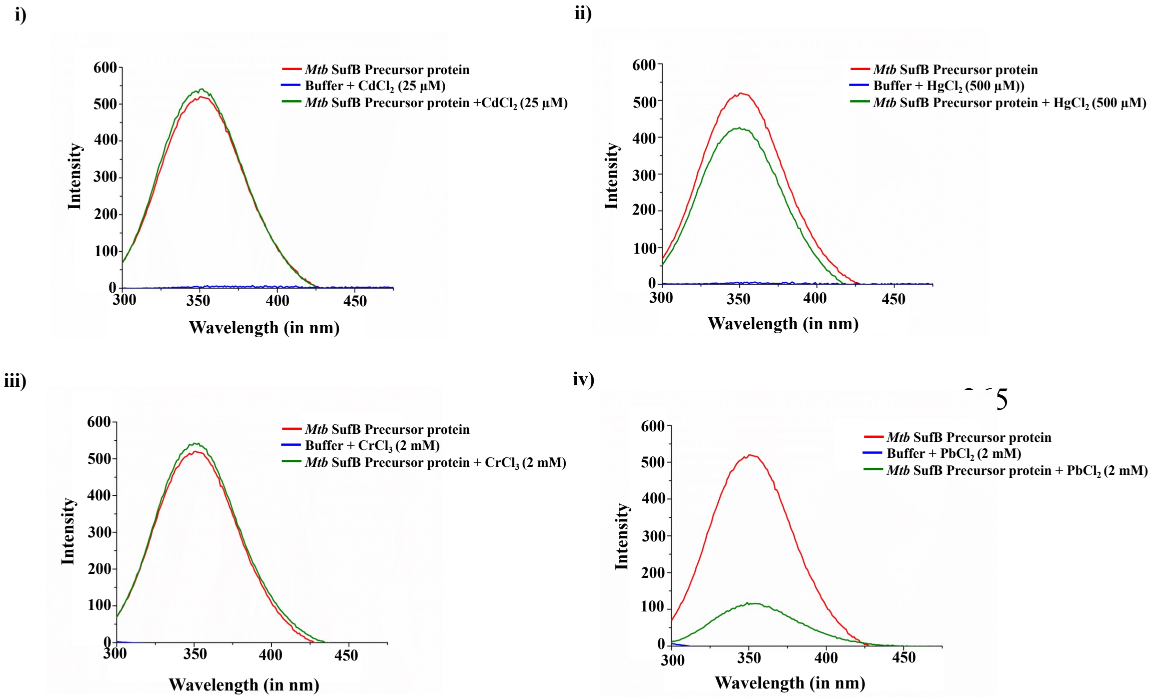

**Figure S7. Tryptophan fluorescence assay to evaluate the interaction between the *Mtb* SufB precursor protein and toxic metal ions.** The protein was incubated with (i) $CdCl_2$ (25 µM), (ii) $HgCl_2$ (500 µM), (iii) $CrCl_3$ (2 mM), and (iv) $PbCl_2$ (2 mM). Emission spectra were recorded from 300 to 600 nm following excitation at 280 nm to assess metal-induced changes in protein fluorescence intensity.

**Quantitative evaluation of *Mtb* H37Ra-based biosensor platform for the detection of toxic metal ions**

Alamar Blue assay is a qualitative as well as a quantitative test for detecting viable microbial cell population. A spectral absorbance measurement was performed to quantitate the presence of toxic metal ions highlighting the efficacy of the designed mycobacterial whole-cell biosensor platform. These assessments further supported the results from Fig. 2, 3, and 4 (main text). Following 24 hr. of incubation with the Alamar Blue reagent, absorbance was recorded from the respective 96-well plates at 570 nm, with 600 nm as a reference wavelength. Subsequently, a standard plot was generated to determine the concentrations of $Cd^{2+}$ and $Hg^{2+}$ in the test samples. The details of the standard plot and its corresponding linear regression analyses are provided in the main text ( *Mtb* H37Ra-based biosensor platform for detection of toxic metal ions; Results and Discussion).

Four unknown samples (Unknown 1, Unknown 2, Unknown 3, and Unknown 4) were prepared with varying concentrations of $CdCl_2$ ions and $HgCl_2$. The concentrations of these samples were taken arbitrarily and then determined via the standard plot. The purpose of this experiment is to validate the utility of standard plot by extrapolating the absorbance of unknown samples and determine the respective $Cd^{2+}$ ions and $Hg^{2+}$ ions concentrations.

*Unknown 1:* The measured absorbance of Unknown 1 was 0.103, which corresponds to a $Cd^{2+}$ concentration of 9.8 micromolar (μM) based on the standard calibration plot whereas the absorbance of Unknown 1 was 0.075, which corresponds to a $Hg^{2+}$ concentration of 26.36 micromolar (μM) based on the standard calibration plot.

*Unknown 2:* The measured absorbance of Unknown 2 was 0.071, which corresponds to a $Cd^{2+}$ concentration of 5.17 micromolar (μM) based on the standard calibration plot whereas the absorbance of Unknown 2 was 0.285, which corresponds to an $Hg^{2+}$ concentration of 90 micromolar (μM) based on the standard calibration plot.

*Unknown 3:* The measured absorbance of Unknown 3 was 0.314, which corresponds to a $Cd^{2+}$ concentration of 40.39 micromolar (μM) based on the standard calibration plot whereas the absorbance of Unknown 3 was 0.198, which corresponds to a $Hg^{2+}$ concentration of 63.63 micromolar (μM) based on the standard calibration plot.

*Unknown 4:* The concentration for Unknown 4 could not be detected as the concentration of $Cd^{2+}$ and $Hg^{2+}$ in this sample was beyond the linear range of the standard calibration plot. To accurately determine the concentration of $Cd^{2+}$ and $Hg^{2+}$ in such oversaturated samples, the sample needs to be diluted and re-measured.

Overall, the experiment successfully validated the standard calibration plot for $Cd^{2+}$ and $Hg^{2+}$ concentrations within the measured range. However, the high concentration in Unknown sample 4 caused a saturation effect, preventing the accurate determination of its $Cd^{2+}$ and $Hg^{2+}$ concentrations.

**Table S1.** The absorbance value of unknown samples and their corresponding concentrations as obtained from the linear regression plot as shown in Fig. 3.

| | **Unknown sample 1** | | **Unknown sample 2** | | **Unknown sample 3** | | **Unknown sample 4** | |
|---|---|---|---|---|---|---|---|---|
| **Toxic metal ions** | **$Cd^{2+}$** | **$Hg^{2+}$** | **$Cd^{2+}$** | **$Hg^{2+}$** | **$Cd^{2+}$** | **$Hg^{2+}$** | **$Cd^{2+}$** | **$Hg^{2+}$** |
| **Absorbance** | 0.103 | 0.075 | 0.071 | 0.285 | 0.314 | 0.198 | Conc. exceeded | Conc. exceeded |
| **Concentration in μM** | 9.8 | 26.36 | 5.17 | 90 | 40.39 | 63.63 | Conc. exceeded | Conc. exceeded |
